# Template-free detection and classification of heterogeneous membrane-bound complexes in cryo-electron tomograms

**DOI:** 10.1101/413484

**Authors:** Antonio Martinez-Sanchez, Zdravko Kochovski, Ulrike Laugks, Johannes Meyer zum Alten Borgloh, Saikat Chakraborty, Stefan Pfeffer, Wolfgang Baumeister, Vladan Lucic

## Abstract

With faithful sample preservation and direct imaging of fully hydrated biological material, cryo-electron tomography (cryo-ET) provides an accurate representation of molecular architecture of cells. However, detection and precise localization of macromolecular complexes within cellular environments is aggravated by the presence of many molecular species and molecular crowding. We developed a template-free image processing procedure for accurate tracing of complex networks of densities in cryo-electron tomograms, a comprehensive and automated detection of heterogeneous membrane-bound complexes and an unsupervised classification. Applying this procedure to tomograms of intact cells and isolated endoplasmic reticulum (ER), we detected and classified small protein complexes like the ER protein translocons, which were not detected by other methods before. This classification provided sufficiently homogeneous particle sets and initial references to allow subsequent de novo subtomogram averaging. Therefore the procedure presented allows a comprehensive detection and a structural analysis of complexes in their native state. In addition, we present structural evidence that different ribosome-free translocon species are present at the ER membrane, determine their 3D structure, and show that they have different localization patterns forming nanodomains.

## Introduction

The cellular environment is characterized by the presence of many different molecular species. Complexes, stable or transient underlie critical cellular functions. Of particular interest are membrane-bound complexes because they are essential for many types of cellular processes, like cell signaling, immune response and synaptic transmission, and are targeted by more than two thirds of all drugs [1].

In cryo-electron tomography (cryo-ET), biological samples are faithfully preserved by rapid freezing, which prevents water crystallization and rearrangements of the biological material. Importantly, samples are imaged in transmission electron microscopy in the same vitrified, fully hydrated state [2, 3]. Therefore, cryo-ET is uniquely suited for high resolution, direct three-dimensional (3D) imaging of unperturbed cellular environments [4, 5].

The potential of cryo-ET to yield cellular maps of molecular complexes is hampered by low signal-to-noise ratio (SNR) in tomograms and the molecular heterogeneity in cells. Because visual detection is limited to large complexes of characteristic shapes [6], image processing methods have been developed to interpret tomograms. In template matching, currently the leading detection method in cryo-ET, a structure of a protein or complex of interest is used to computationally search for similar structures [7, 8, 9, 10, 11, 12]. This approach is particularly suited for complexes that do not form part of larger assemblies and critically depends on the already existing structures of complexes of interest. Automated methods were developed for segmentation of cellular components of particular shape, such as lipid membranes and filaments [13, 14, 15]. Pleomorphic, membrane associated complexes were previously detected by an automated method [16], but their molecular identification remained challenging [17, 18, 16]. Subtomogram averaging can yield 3D densities at a resolution higher than that of individual complexes, but requires biological systems where a complex of interest is present in a large number [19, 20, 21].

### Algorithm 1 Complete procedure

1. Density tracing and particle picking
  - Tracing of density by the Discrete Morse theory based algorithm (DisPerSe)
  - Simplification by topological persistence
  - Spatially embedded graph representations of the EM density
  - Selection of complexes - particle picking
2. General classification
  - Determination of membrane normal vectors
  - Constrained refinement (Relion)
  - Unsupervised classification of rotationally averaged complexes by Affinity propagation
3. Spatial analysis and averaging
  - Standard 3D classification and constrained refinement (Relion)
  - Spatial distribution analysis within or between classes

To allow comprehensive, high resolution processing of cellular cryo-tomograms, we developed a software procedure for template-free detection and unsupervised classification of heterogeneous membrane-bound molecular complexes. It includes methods from other fields that we adapted and further developed, such as the discrete Morse theory based segmentation, affinity propagation (AP) and spatial point processes, as well as custom-made software. The classes obtained were sufficiently homogeneous to allow further processing by standard subtomogram averaging methods [22, 21]. Validations and comparisons with other methods were performed on phantom and real datasets.

## Results

### Procedure overview

Our procedure consists of three major parts (Algorithm 1). First, electron density is traced and complexes are detected in a comprehensive, template-free manner. Then, they are classified into classes containing structurally similar complexes, rendering them suitable for further processing. Finally, the spatial distribution of complexes and their average densities are determined.

### Density tracing and simplification

For an automated tracing of density in cryo-tomograms, we developed a procedure based on DisPerSE, a software package which employs discrete Morse theory originally developed to identify astrophysical structures in 3D images of the large-scale matter distribution in the Universe [23]. In general, Morse theory is used to calculate topological indices (invariants) of a given manifold, while the discrete Morse theory is applied to simplicial complexes [24, 25].

#### Algorithm 2 Simplification by topological persistence

For each pair of connected minima and saddle points (p_i_, s_a_) whose values differ by less than a specified persistence value:

1. Find the other minimum (p_k_) connected to the saddle point s_a_ and connect it to all saddle points connected to p_i_
2. Remove p_i_, s_a_and all their arcs
3. Add ascending manifold associated with minimum p_i_ to the one associated with minimum p_k_
4. Remove arcs associated with saddle points of low density

We used DisPerSE to generate Morse complexes comprising the following manifolds: (i) Greyscale minimum points (termed 0-critical points), (ii) Saddle points that have minima in two and a maximum in one direction (1-critical points), (iii) Arcs that connect minima and saddle points defined as maximum gradient curves between these points, (iv) 3-manifolds associated with minima (ascending 3-manifolds). To better visualize the tracing, we processed a (2D) tomographic slice of a neuronal synapse (Figure 1 A, C). The minima and the arcs visually corresponded well to the distribution of the electron density.

**Figure 1:**
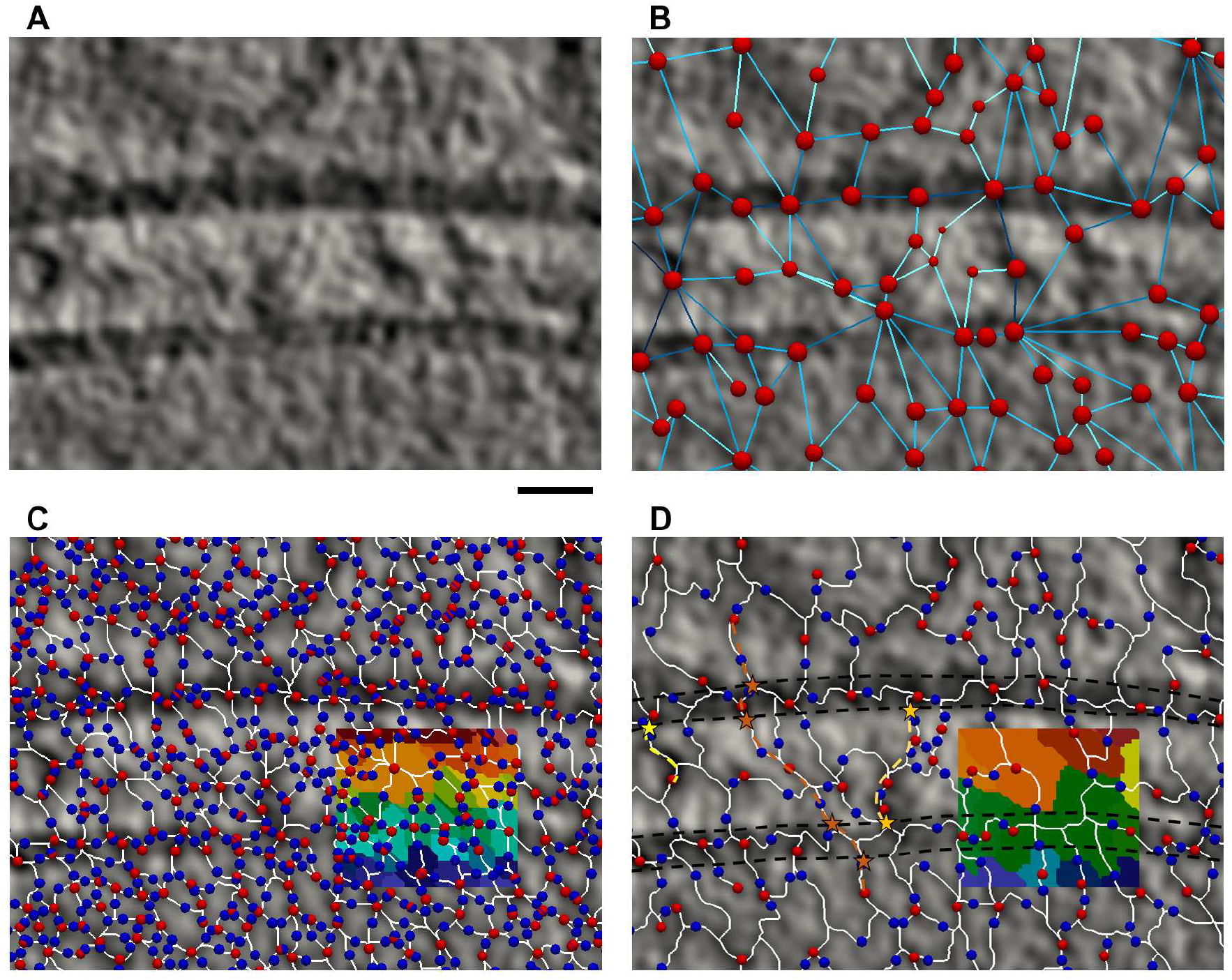
Tracing density at a synapse. A Tomographic slice of 1.37 nm thickness. B Graph representation of the Morse complex shown in D. Note that although graph edges are shown by straight lines, they keep the full information about the arcs they represent shown on D. C Morse complex obtained by application of DisPerSE on the slice shown in A. D Morse complex obtained from C after simplification by topological persistence. B-D Images superimposed on the slice from A. C, D Color insets show ascending 2-manifolds, labeled by different colors. Red circles represent greyscale minima and graph vertices, the blue ones the saddle points, white lines arcs and blue lines graph edges. In B, larger size vertices denote minima of higher density and darker shade edges denote saddle points of higher density. Yellow, orange and brown paths represent possible extracellular presynaptic, trans-cleft and extended trans-cleft complexes. Asterisks are intersection points between the selected paths and membrane faces. Black dashed lines outline synaptic membranes. Scale bar 10 nm.

A high level of noise present in cryo-tomograms causes the detection of many closely spaced local minima that have only slightly smaller values than their neighborhood, resulting in an overly complex structure of the calculated Morse complexes. Because of a highly complex network of densities present in cellular cryo-tomograms, we could not use the Morse complex simplification procedure implemented in DisPerSE. To solve this problem, we implemented a modified version of the simplification by topological persistence [23]. Namely, a saddle point and its adjacent minimum that have similar grayscale values are removed (low persistence pairs), and the affected Morse complex elements are reassigned. (Algorithm 2, Figure S1). This is equivalent to introducing small perturbations in the greyscale values that remove some minima, effectively reducing the contribution of noise.

The simplification by topological persistence resulted in a greatly simplified Morse complex, and a faithful tracing of density (Figure 1D). All together, the choice of manifolds provided by the discrete Morse theory, combined with the custom-made implementation of the simplification procedure, made it possible to accurately trace electron density in cellular cryo-tomograms.

### Graph embedding and detection of complexes (particle picking)

To streamline further processing, we implemented a procedure that converts the information obtained by the density tracing to spatially embedded graphs. Namely, each arc is assigned to a graph edge and its two adjacent minima are assigned to graph vertices associated with the edge (Figure 1B). Vertices and edges keep the information about the spatial location, greyscale density, geometry and the connectivity of the underlying minima and arcs. Also included are external information about the identity of cellular structures, such as the lipid membranes. Therefore, these spatially embedded graphs represent the distribution of the biological material (proteins and lipids) visualized in a tomogram.

They occupy the central part in the software we developed, because they combine precise geometrical, localization and biological information, and allow computationally efficient queries that can extract specific information used to detect individual complexes (particles). Importantly, subgraphs can be selected starting from vertices belonging to, or at a specified distance from a previously defined membrane or another cellular structure. Greyscale values and geometrical information associated with vertices and edges can be used as further constraints. For example, the yellow and the orange star-bound paths in Figure 1D likely represent an extracellular presynaptic membrane-bound and a transcleft complex, respectively, while the brown path shows a putative complex that directly links pre- and postsynaptic intracellular components. Not all selected subgraphs can be directly matched to complexes, even though subgraphs represent densities. The repositioning or elimination of wrongly selected subgraphs is delegated to the subsequent processing steps. Hence, the procedure described so far corresponds to particle picking in the single particle analysis.

### Density tracing validation on phantom data

To validate the Morse theory-based density tracing and the simplification procedures, we created a phantom dataset comprising a rectangular grid having higher density at the intersections, and added variable amounts of Gaussian noise (Figure S2A). Densities in all phantom datasets were detected by applying the discrete Morse theory and the topological simplification as outlined above. Grid intersections and grid bars were taken as the ground truth features for the detection of minima and arcs, respectively. We detected the minima and arcs that matched the ground truth (true positives, TP), did not match the ground truth (false positives, FP), as well as the unmatched ground truth features (false negatives, FN) (Figure S2B). Multiple minima occurring at the same grid intersection were avoided by imposing an exclusion distance between particles.

Numbers of TPs, FPs and TNs were normalized to the total number of the corresponding ground truth features. For SNR above 0.05, the FPs and FNs were below 10% and TP minima was above 90%, however for SNR between 0.05 and 0.1 TP arcs was between 80% and 90% (Figure S2C). To a large extent, this failure to detect some of the ground truth arcs (these constitute FN arcs) was caused by the minima that were not detected (FN minima). This was confirmed by normalizing TP arcs to the total number of ground truth arcs that could be formed given the detected minima (TP arcs corrected in Figure S2C).

### General classification

The application of the procedure described in the previous section to complex cellular systems is expected to yield a set of membrane-bound complexes possessing high compositional and conformational heterogeneity. Therefore, a general classification procedure capable of separating highly heterogeneous complexes into groups (classes) of similar complexes is required for further processing.

Particle (complex) positions are used to calculate the direction of vectors perpendicular to the membrane. These normal vectors specify two of the angles that determine the orientation of the particles, while the third angle is left undetermined. To optimize normal vectors and particle positions, we employ constrained particle refinement. That is, the two Euler angles that define the normal vectors are allowed only small changes around the initial values during the alignment step, while the third angle (around the normal) is not constrained. Furthermore, a high symmetry imposed on the third angle diminishes its importance for the alignment. The initial reference for this refinement can be obtained from the data, by randomizing the third angle and averaging all particles (without alignment). The symmetrization around the normals and the choice of the initial reference reduce the influence of the missing wedge.

The methods chosen for particle classification is the affinity propagation (AP) clustering [26], whereby nodes (particles) exchange information between each other to reach the optimal partitioning. The advantages of the affinity propagation compared to the standard clustering methods are that this algorithm is unsupervised, the number of classes is not specified in advance but data driven and it can handle cases where classes have a very different number of particles.

The success of a clustering procedure critically depends on the manner the clustering distance (similarity) is defined. Here, we represented particles as 2D images obtained by computing particle rotational averages around their normal vectors (illustrated on Figures S6, S3). We defined the distance between two particles as the dot product of their normalized rotational averages. In this way, the 2D averages used for clustering were pre-aligned to each other, and there were no further degrees of freedom that could affect the clustering. On the contrary, if clustering was based on the 3D particle subtomograms, the angle around the normal vector would need to be determined, thus hampering the procedure.

### Validation of general classification

To evaluate the AP classification independent of particle picking and to compare it with two other methods used in the field (K-means and hierarchical clusterings) we generated a test dataset from eight available reference structures of membrane-bound complexes. To make realistic particles we added different amounts of noise, tilted them around the membrane normal, rotated around the normal, translated them along the membrane plane and imposed the missing wedge (see the Methods; Figure S3A).

We classified the test dataset and compared results against the ground truth using three measures. The first two, Fowlkes-Mallows [27] and the Variation of information [28] compare two different classifications directly and are currently the state-of-art methods in the clustering analysis, but are rarely used in cryo-ET [16]. We also made a correspondence between the classes obtained and the ground truth classes, based on the number of reference structure elements within each class, from which we determine the F_1_ measure (see the Methods).

Our data for all three metrics shows that the AP classification is weakly affected by changes in SNR, displacement range and the number of reference structures (Figure S4A-C). The input preference parameter that has to be specified for AP classification only weakly influenced the scores. It showed the best results for the input preference parameter values between −6 and −3 and good results between −10 and −2, which defines the range of preference parameter values suitable for cryo-tomograms. In addition, the high similarity between the AP class representatives (called examplars) and the ground truth class averages further confirms the strength of the AP classification (Figure S3B).

K-means and hierarchical clustering methods require the number of classes as an input parameter, but this is usually not known in advance. Our data showed that for the optimally chosen number of clusters, which is not possible without external information, the performance of K-means and hierarchical clusterings was similar to the AP using the default input preference values. However, it was drastically reduced for realistic cases when non-optimal number of classes is chosen (Figure 2A). The results obtained using r-weighted averages (see the Methods) brought the same conclusions, but the scores improved (Figure S4D). Therefore, the distinctive advantage of the AP classification is that it yields optimal results without requiring external prior information.

**Figure 2:**
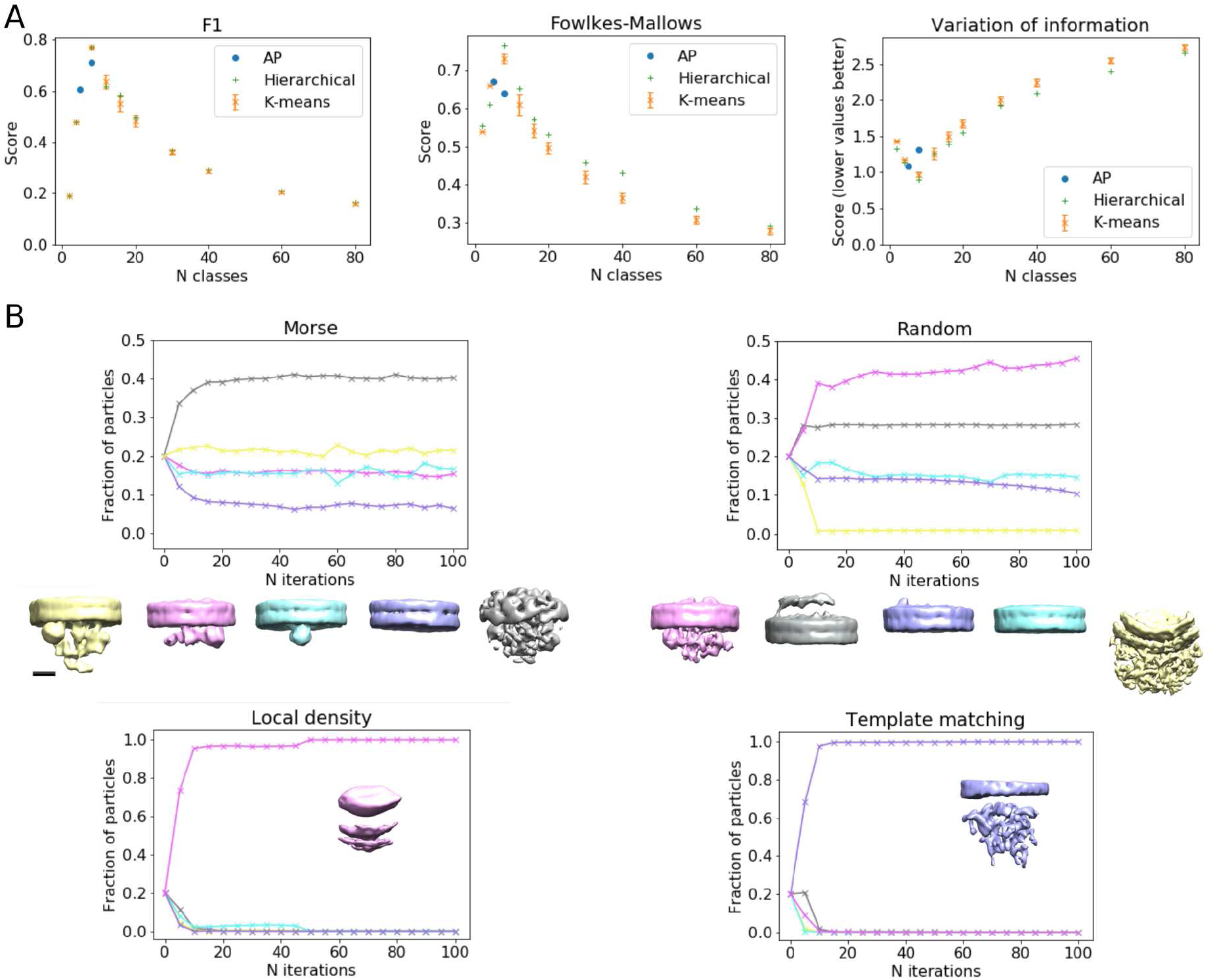
Method validations. A Comparison of the AP with K-means and hierarchical clusterings. F_1_(left), Fowlkes-Mallows (middle) and Variation of information (right). Note that higher values of the Fowlkes-Mallows and F_1_ measures, but lower values of the Variation of information signify better agreement. The AP clustering results are shown for the preference parameter taking all integer values between −9 and −3 (several data points overlap). For K-means clustering the data points denote means and the error bars the standard deviation obtained from 200 simulations. B Validation of particle picking, graphs showing the number of particles per class are shown together with the class averages. Morse picking (up, left), random (up, right), local density (bottom left) and template matching (bottom right). Scale bar 5 nm.

### Detection and classification on microsomal membranes

To test the particle localization and general classification methods introduced above, we used a subset (26%) of previously analyzed cryo-ET data depicting canine pancreatic microsomes [29]. This work established the basic architecture of the translocon complex and structure of its constituents: the Sec61 protein-conducting channel, the translocon-associated protein complex (TRAP) and the Oligosaccharyl-transferase complex (OST) [30, 31].

We traced biological material as explained above (Figure 3A, Video 1). Particles at the cytoplasmic and lumenal faces of the ER membrane were located independently of each other, based on minima that satisfied certain geometrical constraints (see the Methods). Particle positions and the membrane normal vectors were optimized by the constrained refinement with C10 symmetrization, yielding average densities having a well-positioned density and a resolved lipid bilayer (Figure 3B).

**Figure 3:**
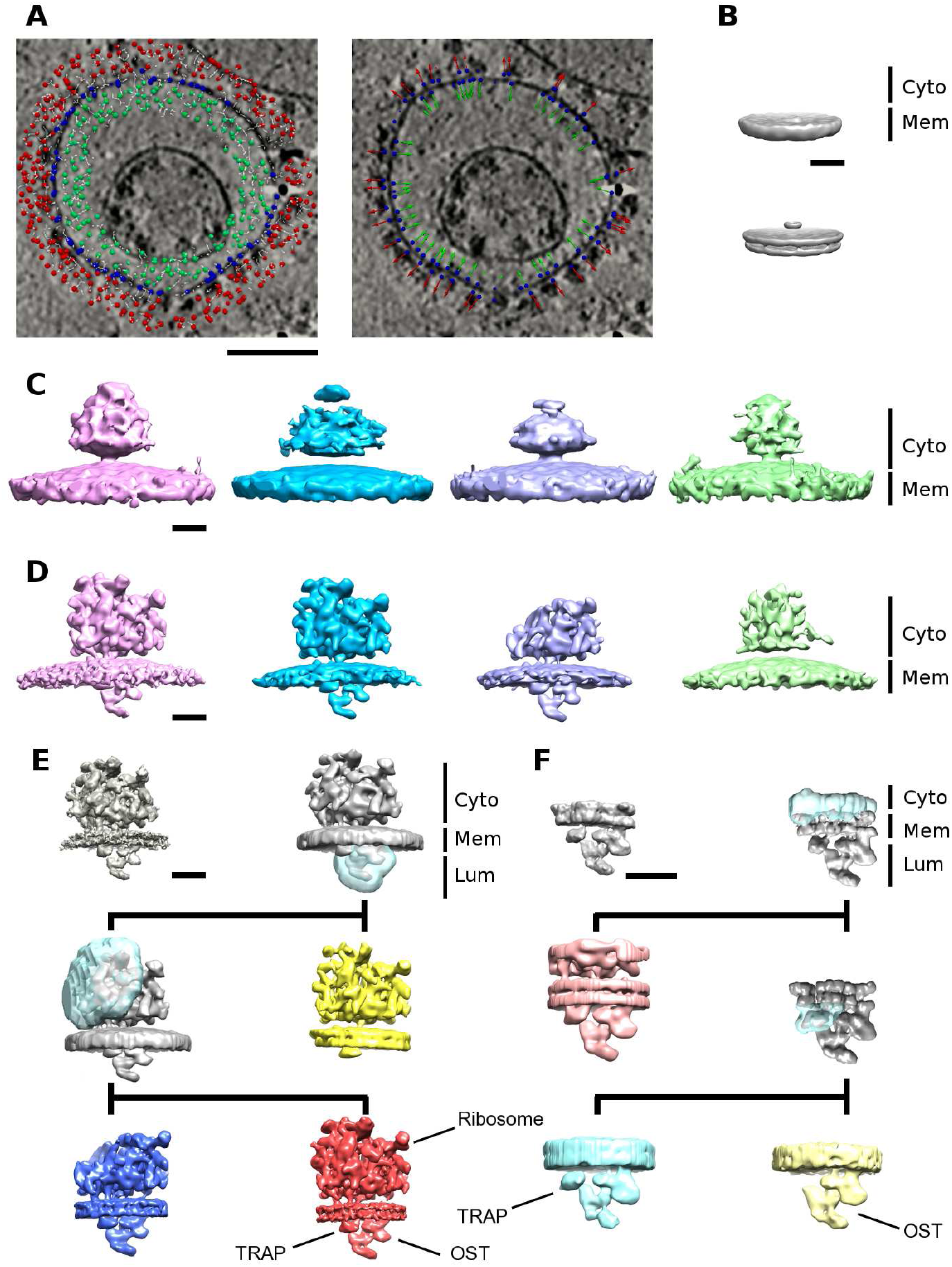
Processing of ER membrane-associated complexes. A Density tracing. Density minima are shown as small spheres (left) and particles as spheres with arrows representing the associated membrane normal vectors (right). Position of the spheres is color-coded: Red cytosol; blue membrane; green lumen. B Average of all cytosolic particles obtained without alignment (above) and with constrained refinement with ClO symmetry (below). C, 3D class averages of the representative affinity propagation classes of cytosolic particles (ribosomes), obtained without alignment. D Refinement of the same classes shown in (C) obtained using the averages shown in (C) as initial models. E 3D classification and refinement of cytosolic particles (ribosomes). Densities of ribosomes bound to the fully assembled (red) and partial translocon complex (yellow), as well as the large ribosomal subunit bound to the fully assembled translocon (blue) are shown. F 3D classification and refinement of lumenal particles (translocon). Densities of the ribosome-translocon complex (light red), the ribosome-free fully assembled translocon (light blue) and the non translocon associated individual OST complex (light yellow) are shown. In both **(E)** and **(F)** initial references obtained from the AP classes (top row, left) and densities obtained by the first classification of all particles (top row, right) are shown together with the classes obtained by the second (middle row) and the third round of classification (bottom row). Transparent blue regions correspond to the masks used for classification. Refined and post-processed densities are shown in color. ‘Cyto’, ‘Mem’ and ‘Lum’ denote cytosolic, membrane and lumenal regions, respectively. Scale bars A 100 nm, B-F 10 nm.

Classification of cytosolic particles by AP yielded more than 100 classes (Figure S5). Constrained refinement of these classes, using internal initial references, showed different species of ribosome-translocon complexes (Figure 3C, D). The best class that contained both cytosolic and lumenal density was used to generate an initial reference for further processing (Figure S6).

All cytosolic particles were subjected to three rounds of 3D classification. In the first variant of data processing (termed “bulk cleaning”), all particles were subjected together to a first 3D classification to remove suboptimal particles (Figure S6, dotted line). The second 3D classification round focused on the lumenal segments and the third on the small ribosomal subunit, resulting in structures comparable to those previously reported (Figure S7) [30, 29]. These included fully assembled ribosomes bound to the fully assembled and partial translocon complexes, as well as ribosomal large subunits. This confirms that sufficiently homogeneous particle sets were generated by our procedure, which could be further processed by standard external reference-free subtomogram classification and averaging.

In the second variant of data processing (termed “AP cleaning”), the first 3D classification was applied to each AP class separately (Figure S6, dashed line) and the other two 3D classification rounds were performed as before. This variant yielded the same ribosomal species, but TRAP was better resolved in the partial translocon complex class, the number of particles was increased and the resolution obtained was slightly higher (18 Å and 22 Å for fully assembled ribosomes bound to the fully assembled and partial translocon complexes respectively, and 21 Å for ribosomal large subunits) (Figure 3E). Therefore, in addition to providing an internal initial reference, the affinity propagation classification thus contains information that can be exploited by subsequent processing.

Lumenal particles were classified by affinity propagation and the best out of 100 classes was refined to yield a 3D average of the translocon (Figure S5B). The bulk cleaning variant, using the translocon average as the initial reference, yielded a well resolved ribosome-translocon class and classes representing two different ribosome-free translocon states (Figure 3F). Among the particles that contained a defined lumenal density, 15% had an associated ribosome and thus corresponded to the ribosome-translocon complex. Among the ribosome-free complexes, 68% corresponded to fully assembled translocon (TRAP, OST and Sec61) and 17% likely represented individual OST complexes. These structures were resolved to 22 Å, 14 Å and 16 Å, respectively (Figure S5D). Importantly, without the initial reference generated from affinity propagation classification, the same 3D classification procedure failed.

Therefore, our Morse theory-based detection is capable of picking small complexes, like the translocon or even smaller individual OST complexes (≈200 kDa lumenal mass). The unsupervised classification by affinity propagation was instrumental to carry the processing to a level where the external reference-free standard 3D classification and refinement procedures could be used.

### Validation of density tracing and particle picking on microsomal membranes

Next, we compared the performance of our Morse theory-based density tracing and particle picking method with the standard methods used in the field: template matching, local density detection (after low-pass filtering) and random picking. In all cases we analyzed a subset of tomograms used in the previous section and picked particles on the lumenal side of the microsomes, up to 20 nm to the membrane. Template-matching was done using the ribosome-free translocon average density obtained in the previous section (Figure 3F).

The picked particles were subjected to a 3D constrained refinement with a high symmetry, followed by 3D classification that used the same ribosome-free translocon density as the initial reference. This procedure differs from the one presented in the previous section in that it skips the general classification, but relies on an external initial reference. In all cases, the classification converged to a stable number of particles after 15-20 iterations (Figure 2B). All five classes from Morse-picked particles contained a significant number of particles. One class represented the translocon complex, while another two classes likely represented two other small membrane-bound complexes. Among the last two classes, one probably contained false positive picks because the average showed just a well-resolved membrane bilayer, while the other class likely contained a mixture of other complexes. This shows that small complexes can be reliably detected by Morse picking.

Four classes of random-picked particles contained a significant number of particles, but none of them yielded recognizable densities. The local density detection and template matching yielded particles that all converged to one class and produced poor averages. Therefore, among the particle picking methods tested, the Morse method was the only one that successfully detected small complexes in general and the translocon complex in particular.

### Detection and classification in situ

We applied our procedure for the detection and classification of membrane-bound complexes to tomograms of intact P19 embryonic carcinoma cells. After the Morse-based density tracing, particles were picked in the vicinity of endoplasmic reticulum (ER) membranes. Because the orientation of the membranes could not be always determined and the cellular interior is densely populated, we expanded the general classification to include three rounds of AP classification. The first AP classification served to clean the particle set and to group other classes into datasets defined by the following densities: large densities located on the same side of the membrane where particles were picked, large densities on the side opposite from the particles and the small densities (Figure S8). After the subsequent two AP classification rounds, class averages were obtained for selected classes (Figure S9).

As expected, ribosomes were prominent in the AP classes derived from the large densities on the large density on particle side dataset (Figures 4, S9A). Furthermore, we obtained other class averages that showed a distinct morphology. One of them we tentatively assigned to the Inositol trisphosphate (IP3) receptor and two other to various assembly states of MHC-I peptide loading complex (PLC), based on the similarity with densities previously obtained by single particle analysis [32, 33].

**Figure 4:**
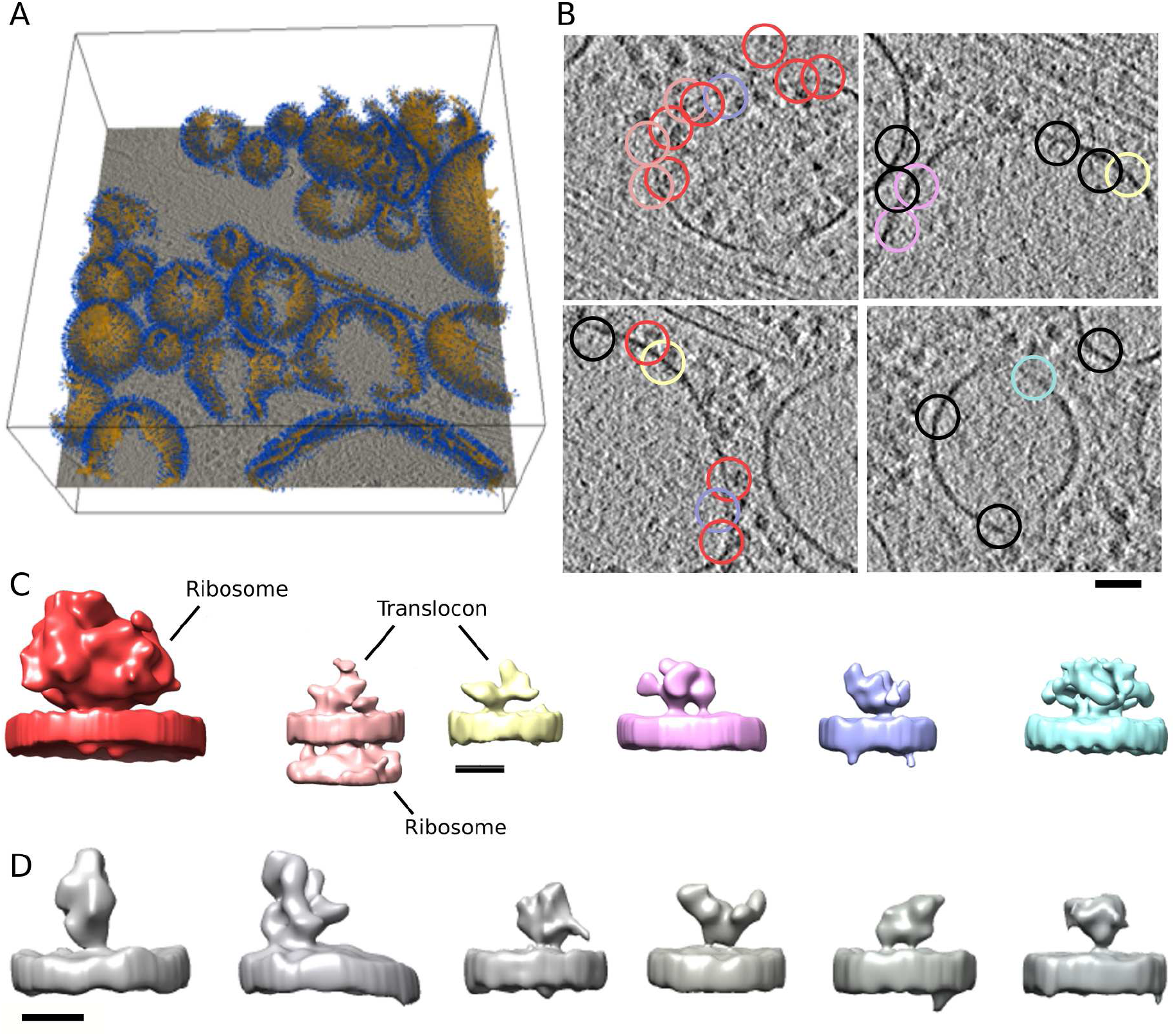
Complexes resolved in situ. A Tomographic slice. Superposed are segmented internal membranes (ochre-yellow) and picked particles including the normals (blue). B Tomographic slices showing x and y positions of some particles classified as complexes shown in C (colors match). The exact particle positions can be on tomographic slices up to 20 nm away from the shown slices. Scale bar 50 nm. C Average densities, from left to right: membrane-associated ribosome (red), ribosome-associated translocon (light red), ribosome-free translocon (yellow), two putative PLC complexes (magenta and purple), putative IP3 receptor (blue). Scale bar 10 nm. D Selection of the other class averages. Scale bar 10 nm.

The ribosome-translocon complex was detected in the dataset containing large densities on the opposite side of the membrane (Figures 4, S9B). It is important to note that the Morse based procedure picked the translocon complex directly, while the large densities (ribosomes) seen in the AP class averages arose during the classification.

Importantly, we obtained an AP class average from the small densities dataset that was visually consistent with our ribosome-free full translocon and OST complexes from the microsomal data, as well as with the ribosome-bound translocon from the in situ data (Figures 4, S9C, 3F). We also generated several other AP averages showing small densities, the identification of which would require a more in-depth analysis. Inspection of these classes showed homogeneous sets of particles, confirming the validity of the classification.

Particles randomly chosen from the small density dataset after the first, and those chosen after the second AP classification were refined. Both resulting averages showed only the membrane, confirming the importance of our AP - based general classification approach. All together, our procedure is applicable to cryo-tomograms of intact cellular interior and allows the detection of even small ER membrane-bound complexes and their separation into classes that are sufficiently homogeneous for standard processing.

### Spatial distribution analysis

Methods available for the analysis of spatial point processes can provide further information about the distribution and clustering of complexes and assist with classification. Specifically, monovariate distribution functions analyze a single class. Among these, the first order functions are based on the distance to the closest point, either from other points (nearest neighbor distribution) or from arbitrary locations (spherical contact function), or both (J-function) [34]. A more detailed description is obtained by Ripley’s second order functions, which evaluate the distribution at different length scales, by considering distances between all pairs of points [35, 36]. Furthermore, bivariate versions of the nearest neighbor and Ripley’s functions characterize colocalization and co-clustering of particles between two classes.

To assess the statistical significance, the above functions are evaluated with respect to the random distribution (null hypothesis). Due to the restricted and irregular shape of the region where the particles are located, analytical models cannot be used. Instead, many random point distributions need to be generated within the particle region. To this end, we implemented a Monte-Carlo method that generates random distributions of a specified number of particles within an arbitrary space (see Methods).

### Spatial organization of microsomal complexes

As an example of using spatial point distribution methods to address biological questions, we investigated the spatial organization of the microsomal particle classes obtained above. Upon visual inspection, some classes showed distinct distributions (Figure 5A, Video 2). In order to quantitatively determine whether the complexes were clustered and at which length scales, we calculated the univariate Ripley’s function for the particle classes and compared them with results obtained for simulated particles.

**Figure 5:**
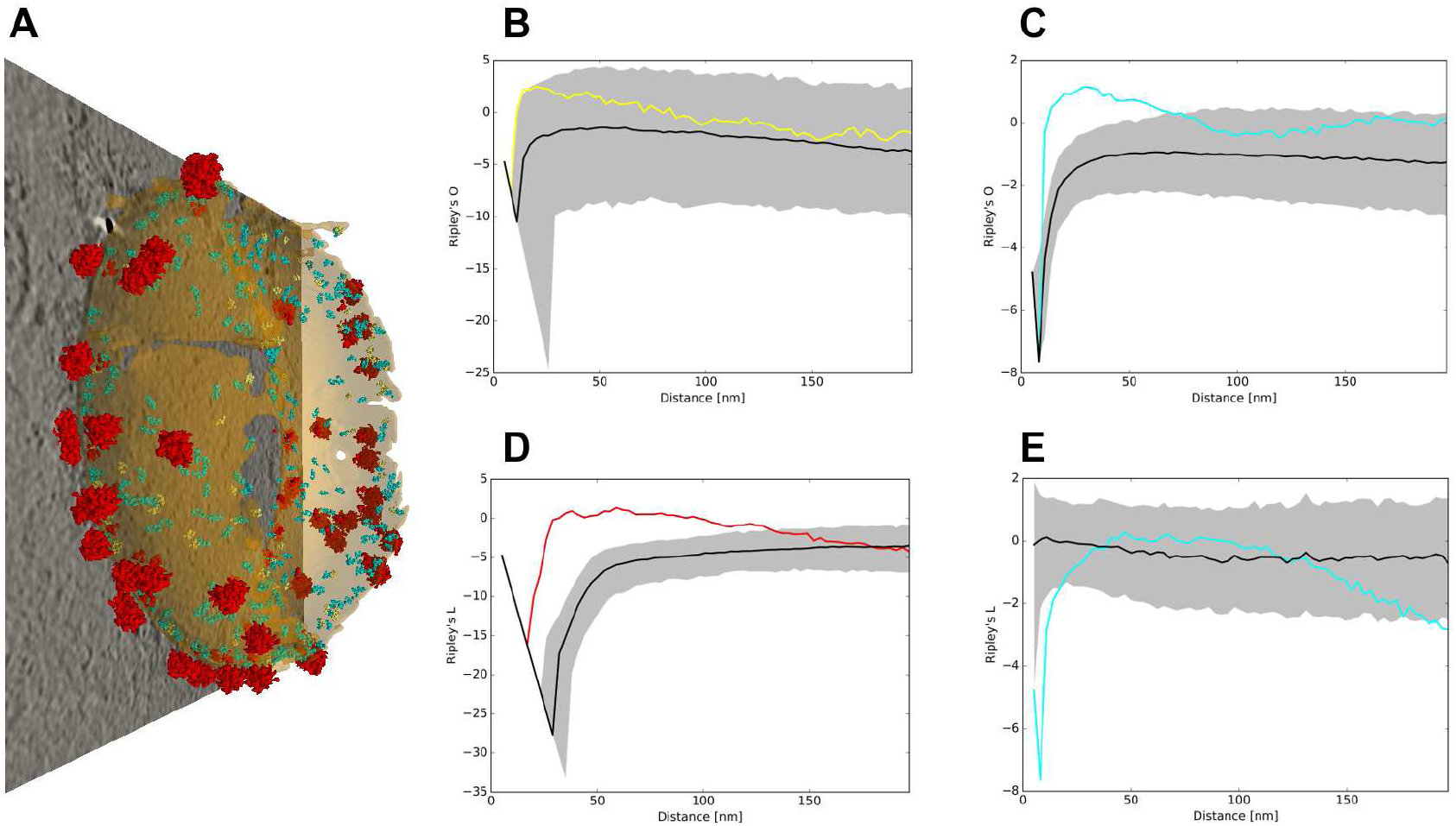
Spatial distribution of microsomal particles. A Distribution of the particles is shown on one microsome (all three cytosolic ribosome classes red, ribosome-free full translocon: cyan, non-translocon associated OST: yellow). B-D Univariate Ripley’s **L** function of the three classes, the same color code as in A. E Bivariate Ripley’s **L** function between the cytosolic ribosome classes and the ribosome-free translocon complex. In B-E Black lines show the median of the Ripley’s **L** function for a set of random particle distributions (≈1200) and the gray areas represent regions of p>0.05 confidence.

The ribosome-free translocon complex showed a significant clustering at length scales from about 8 nm to more than 50 nm, while the non-translocon associated OST was borderline significant (at p=0.05 level) at 10-20 nm in respect to the random distribution (Figure 5B,C). We confirmed these results by using the first order functions: the nearest neighborhood, spherical contact distribution and J-functions all showed significant clustering distribution of the ribosome-free translocon complexes (Figure S10).

As expected, the distribution of ribosomes, comprising all three final classes of the cytosolic particles (Figure 3E), also showed significant clustering (Figure 5D) likely induced by polyribosome formation. In addition, using the bivariate Ripley’s L function, we did not detect a significant colocalization between ribosomes and ribosome-free translocon complexes (Figure 5E, F). These examples show that spatial point process methods allow a quantitative characterization of the organization of molecular complexes.

## Discussion

The mining of the rich information content of cellular cryo-electron tomograms is hindered by by the crowded nature of cells populated by many different molecular species. To overcome this problem, we created a template-free procedure for detection and classification of pleomorphic membrane-bound molecular complexes visualized in cellular cryo-tomograms.

In order to trace density in cryo-electron tomograms, we adapted an existing discrete Morse theory-based procedure and modified the topological persistence simplification [23]. Applications on both phantom and biological data showed accurate tracing of densities at low signal-to-noise conditions characteristic of cryo-tomograms. We developed software to convert the tracing data into spatially embedded graphs that incorporate geometrical, connectivity and greyscale information, which allows automated particle localization. While this Morse theory-based particle picking does not require the presence of membranes, here we included this information to determine position and orientation of particles in respect to local membranes. Direct comparison on real cryo-tomograms showed that our Morse theory-based picking performs much better that other commonly used methods and that it was the only one that could detect small complexes.

Because this rather comprehensive particle picking yields highly heterogeneous particles, may of which do not represent complexes, their structural classification is a challenging task. We found that the combination of unsupervised classification by the AP clustering [26] and the use of 2D particle rotation averages around the membrane normals allows an efficient and successful classification. While the AP classification was previously shown to be superior to K-means and hierarchical clusterings, they performed similarly in our tests when the correct number of classes was set as a parameter to K-means and hierarchical clusterings. However, a distinctive advantage of AP is that it determined the optimal number of classes from the data, without an external input.

Applying our procedure on a dataset depicting isolated microsomes [29], we obtained 3D densities of ribosome-independent translocons, small ER membrane-resident complexes with domains projecting into the lumenal side of microsomes (≈260 kDa total lumenal mass) and even smaller individual OST (≈200 kDa lumenal mass) complexes. These were not previously detected by template-matching on the same dataset. In addition, we obtained 3D densities of different species of ribosomes, consistent with previous template-matching approaches [30, 29]. Small complexes were detected in cryo tomograms before, but not in heterogeneous, cellular systems [37]. All together, these results show the superiority of our template-free procedure.

Furthermore, the application of our procedure on complexes imaged in situ resulted in average densities of several small complexes, including the ribosome-free translocon, medium-sized complexes, as well as ribosomes. This data was consistent with our results from microsomal membranes. The generally lower resolution of the in situ data can be attributed to a more dense and heterogeneous cellular environment and to technical differences between these datasets.

These results demonstrate that the procedure presented here can be applied to small complexes, in their native sate, that were beyond the reach of template-matching. It is not limited by the availability of high resolution structures and, unlike template-based approaches, does not introduce an initial bias that might affect subtomogram classification and averaging. Furthermore, being template-free, our procedure can localize heterogeneous complexes, or complexes that adopt different compositional and conformational states. In both applications, the unsupervised classification by AP provided particle sets that were sufficiently homogeneous for the subsequent standard 3D classification and refinement procedures, as well as internal, data-driven initial references, Together, these results allowed us to determine de novo average 3D densities of several complexes.

The rotational averaging of membrane-bound complexes eliminates the only unknown rotational orientation. Consequently, it removes the problem of incorrect alignment that can hinder classification and diminishes the effect of the missing wedge, while keeping sufficient structural information for classification. The combination of these may be the reason why we successfully applied the AP classification on large datasets (more than 170 000 particles). Because the ability to determine normal vectors is the only membrane-related requirement, our procedure can be applied to complexes attached to any cellular membrane and also larger structures such as the cytoskeleton. The previous template-free approaches, based on automated pattern mining, deep learning and the difference of Gaussians picking methods, were successful only on large complexes, and did not reach resolution comparable to ours, neither on simulated nor on real datasets [38, 39, 40, 15]. Some of these use 3D rotation-invariant properties for classification, a promising method for further developments, but even in the most recent work membrane-attached complexes could not be detected [40]. This argues that processing membrane-bound and membrane-independent complexes are two distinct problems that may require different approaches. Because of the major importance of membrane-bound complexes for biochemical signaling pathways, such as those involved in the immune response, development or synaptic transmission, as well as their dominant role in drug development [1], the procedure described here may be applied to a wide range of fundamental biological processes.

The results obtained from the microsomal data provide structural proof for the presence of ribosome-free translocon complexes in the ER membrane that either await binding of ribosome-nascent chain complexes for co-translational protein transport and membrane insertion, or are engaged in post-translational processes. Notably, the majority of these ribosome-free translocon complexes already comprise all constituents known to be present in the ribosome-associated translocon, arguing against a step-wise assembly of the translocon complex on the ribosome. In metazoans, the STT3A type OST complex is stably integrated into the translocon complex for co-translational glycosylation of nascent proteins, while the STT3B type OST complex is excluded from the translocon and takes care of glycosylation sites skipped by STT3A [41, 42]. Thus, the individual, not translocon-associated OST complexes we localized in the ER membrane likely correspond to STT3B type OST complexes.

Finally, we implemented and adapted the first and the second order spatial distribution functions to characterize point-particle patterns in regions of arbitrary shape. A clustering method based on Voronoi tessellation was recently developed for the detection of large point clusters on a dense background present in super-resolution fluorescence data [43]. This is substantially different from our system, where points (particles) are less numerous but more significant because they were previously selected by stringent classification steps. Using the spatial distribution functions, we observed a significant clustering of ribosome-free translocon complexes on microsomal membranes. This suggests the presence of nanodomains in the ER membrane for post-translational protein transport and membrane insertion, established by direct or indirect interactions between these complexes.

In conclusion, we showed that the template-free procedure presented here is uniquely suited to accurately trace density in cryo-electron tomograms, localize and classify heterogeneous membrane-bound molecular complexes, and provide initial references for 3D refinement. Therefore, it extends the applicability of cryo-ET to small and heterogeneous membrane-bound molecular complexes in their native state and makes possible large-scale, non-invasive detection, localization and de novo structure determination of molecular complexes in situ.

## Methods

### Synaptosomal preparation

Cerebrocortical synaptosomes were extracted from 6–8 week old male Wistar rats as described previously [44, 45, 18] in accordance with the procedures accepted by the Max Planck Institute for Biochemistry. In brief, anesthetized animals were sacrificed, and the cortex was extracted and homogenized in homogenization buffer (HB; 0.32 M sucrose, 50 mM EDTA, 20 mM DTT, and one tablet of Complete mini EDTA-free protease inhibitor cocktail (Roche; 10 ml, pH 7.4) with up to seven strokes at 700 rpm in a Teflon glass homogenizer. The homogenate was centrifuged for 2 min at 2000 g, and the pellet was resuspended in HB and centrifuged for another 2 min at 2 000 g. Supernatants from both centrifugations were combined and centrifuged for 12 min at 9500 g. The pellet was resuspended in HB and loaded onto a three-step Percoll gradient (3%, 10%, and 23%; Sigma-Aldrich) in HB without protease inhibitor cocktail. The gradients were spun for 6 min at 25 000 g, and the material accumulated at the 10/23% interface was recovered and diluted to a final volume of 100 ml in Hepes-buffered medium (HBM; 140 mM NaCl, 5 mM KCl, 5 mM NaHCO3, 1.2 mM Na2HPO4, 1 mM MgCl2, 10 mM glucose, and 10 mM Hepes, pH 7.4). Percoll was removed by an additional washing step with HBM by centrifugation for 10 min at 22 000 g, and the pellet was resuspended in HBM and immediately used in the experiments. All steps were performed at 4 °C.

### Preparation of P19 cells

Murine P19 cells were cultured in alpha-MEM containing nucleosides supplemented with 10% (v/v) FBS, 100 mg/mL each of penicillin and streptomycin at 37°C with 5% CO_2_. Cells were cultivated on Gold Quantifoil grids (R2/1, Au 200 mesh grid, Quantifoil Micro Tools, Jena, Germany). Additional carbon (20–25 nm) was deposited on the film side of the grids in a carbon evaporator (MED 020, BAL-TEC) and plasma cleaned for 45 s prior to their use. Next, grids were sterilized under UV irradiation for 30 min and immersed in culture medium in a CO_2_ incubator for 30 min. Cells were detached from cell culture flasks using 0.05% trypsin-EDTA, seeded on 4-6 pre-treated Quantifoil grids in 35-mm dishes and kept in an incubator overnight to allow adhesion.

### Cryo-ET of synaptosomes and P19 cells

For vitrification, a 3-μl drop of 10-nm colloidal gold (Sigma-Aldrich) was deposited on plasma-cleaned, holey carbon copper EM grids (Quantifoil) and allowed to dry. A 3-μl drop of synaptosomes was placed onto the grid, blotted with filter paper (GE Healthcare), and plunged into liquid ethane.

Grids with P19 cells were blotted from the reverse side by placing a Teflon sheet on the front and vitrified by plunging into a liquid ethane/propane mixture at liquid nitrogen temperature using a Vitrobot Mark 4 (FEI Company, Eindhoven, Netherlands). The Vitrobot was set to 37°C, 90 % humidity, blot force 10, blot time of 10 s and 2 s drain time.

Tilt series were collected under a low dose acquisition scheme using SerialEM [46, 47] on Titan Krios [FEI] equipped with a field emission gun operated at 300 kV, with a post-GIF energy filter (Gatan) operated in the zero-loss mode and with a computerized cryostage designed to maintain the specimen temperature at *<*-150°C. Images were recorded on a direct electron detector device (K2 Summit operated in the counting mode). Tilt series were typically recorded from −60° to 60° with a 2° angular increment. Pixel size was 0.34 nm at the specimen level. Volta phase-plate with nominal defocus of −1 μm [48] was used. The total dose was kept at 60-100 e^−^/Å^2^. Individual frames were aligned using Motioncor2 [49] and dose filtered [50] (P19 cells only). Tilt series were aligned using gold beads as fiducial markers (synaptosomes) or performed by patch-tracking (P19 cells) and 3D reconstructions were obtained by weighted back projection (WBP) using Imod [51]. During reconstruction, the projections were binned once (final voxel size of 0.68 nm) and low pass filtered at the post-binning Nyquist frequency (synaptosomes only).

### Density tracing and particle picking

Electron densities were traced using DisPerSE software package which is based on discrete Morse theory (see [23] for outline of DisPerSE, [25] for a more rigorous presentation of discrete Morse theory and [52] more generally for mathematical topology). Briefly, in Morse theory, a Morse function is defined on a *n*-dimensional manifold. A critical point of the Morse function (points where gradient is 0) that has order *k* has minima in *n-k* and maxima in *k* directions. Gradient paths starting and ending a critical point of order *k* define ascending and descending manifolds of dimensions *n-k* and *k*, respectively. These include local minima, maximum gradient paths connecting critical points and 3D catchment basins. Intersections of ascending and descending manifolds determine Morse-Small cells of dimension *0* to *n*, which partition the space according to the gradient paths and provide connectivity information between critical points. Importantly, 1-critical points are always connected by arcs to two minima. In our case, 3D tomogram greyscale values were used to define a Morse function. Of particular interest are 0-critical points (local minima, Morse-Small *0*-cells), the Morse-Small *1*-cells, that is maximum gradient arcs that connect *0* and *1*-critical points (minima and saddle points that have minima in two and a maximum in one direction). In other words, these trace local density maxima and the “most dense” paths between the densities, thus providing a network that represents EM density. The discrete Morse theory is defined in a similar manner, except that a Morse function is defined on a simplical complex (instead on a manifold), in our case the 3D voxel-based Cartesian grid. The Morse function is then used to determine critical *k*-dimensional simplices, ascending and descending manifolds, and discrete Morse-Small cells.

We modified the topological persistence simplification method and implemented it in PySeg package (Algorithm 2). The procedure first removes the pairs consisting of a minimum and a connected saddle point whose greyscale values differ by an amount smaller than a specified persistence threshold. Then, the arcs and ascending manifolds related to removed points are reassigned. Because this procedure may leave multiple arcs linking the same pair of minima, arcs associated with low-density saddle points are removed. To determine the persistence thresholds, we run multiple simplifications to find the value that gives a specified surface density of minima on membranes. In this way, the density detection is normalized across tomograms. In the same way, (volume) density of minima could be used, so this method is independent of membranes. Furthermore, because the minima are counted, this normalization depends on the total density rather than on the number of complexes of interest.

Density tracing information (specified in the form of Morse complexes) is converted to spatially embedded graphs in the following way. Each arc is assigned to a graph edge and its two adjacent minima are assigned to graph vertices associated with the edge. Vertices and edges keep the greyscale density and a precise location of the underlying minima and arcs. Furthermore, geometrical information, such as the Euclidean and the geodesic lengths of arcs are associated with edges. Additionally, these graphs may also contain external information provided by segmentation of large cellular structures, such as lipid membranes, organelles or cytoskeleton.

Creation of spatially embedded graphs from Morse complexes and their manipulation was implemented in PySeg. For some of the standard graph tasks, the graph-tool library was used [53]. Methods to query properties associated with graph vertices, and edges and methods to extract particles were also implemented in PySeg. All together, representing density by spatially embedded graphs significantly increased the computational efficiency.

### General classification

The general classification was done using the AP clustering algorithm [26]. In short, the algorithm separates elements (here particles) in clusters and determines the representative of each class (“exemplar”). It is based on the iterative calculation of two properties for each pair of elements: the responsibility quantifies how well suited is the first element to be the cluster representative of the second element, while the availability shows how appropriate it is for the first element to have the second element as the representative of its class. In general, initial preferences of each element to be a cluster representative are specified as input parameters. In all our applications, these preferences are taken to be the same for all elements, thus leaving only one parameter. This parameter influences the number of clusters found (the larger the preference the more clusters), but it is not possible to tune the value of the preference so that the classification yields a specified number of classes.

For classification by AP, particles were rotationally averaged around membrane normals by computing mean greyscale values of 2 pixel-wide rings around the normal vectors and the resulting rotational averages were normalized to a density mean of 0 and standard deviation of 1. As an alternative method, weighting the rotational averages by the distance to the axis (r-weighting) emphasizes off-axis elements. For the classification, the similarity between two particles was defined as the dot product between the rotational averages Unless noted otherwise, the masks used for the AP classification cylindrical.

Rotational averaging of subtomograms around the membrane normal vectors and an interface for the scikit-learn [54] implementation of the affinity propagation algorithm are provided in PySeg.

### Density tracing in phantom data

The phantom dataset contained a 6×6×3 grid with a variable amount of Gaussian noise (SNR between 0.005 and 5). For each SNR, 10 datasets were generated. The size of intersections was 2×2×2 voxels and of grid bars 8×2×2 voxels. These datasets were processed in 3D using our procedure. The persistence threshold was set so that the number of minima was 20% higher than the number of grid intersections. The low-density saddle points were removed to obtain 20% more arcs than grid bars, resulting in a higher ratio of arcs to minima (2.3) than the default (2.0), which better captured the high connectivity of the phantom grid. TPs, FPs and FNs were normalized to the total number of ground truth features (grid intersections and arcs). In order to remove the influence of the detection of minima on arc detection, we also normalized the TP arcs to the corrected number of ground truth arcs, that is the number of arcs that could be formed given only the detected minima.

### Validation of general classification

To generate a realistic set of particles, eight available high resolution structures of membrane-bound complexes of different size and shape (4UQJ, 4PE5, 5IDE, 5GJV, 5KXI, 5TJ6, 5TQQ and 5VAI [55, 56, 57, 58, 59, 60, 61, 62]) were selected as the reference structures. First, they were low-pass filtered at 0.524 nm, random noise was added to SNRs of 0.05, 0.01 and 0.005, missing wedge of ±30°was imposed. Then, particles were randomly tilted by 0 - 20° with respect to the membrane normal, rotated around the normal (full range) and displaced along the membrane 0-9 pixels. To test specific parameters, the following test datasets were created from these particles: (i) one for each SNR (Figure S4A), (ii) one for each displacement range (0-3, 0-6 and 0-9 pixels; Figure S4B) at 0.01 SNR, (iii) derived from 4-8 structures at 0.01 SNR (Figure S4C). (iv) For the comparison between the clustering methods, all parameters were randomized, including the number of particles obtained from each reference structure (800 particles in total), and SNR was between 0.01 and 0.005 (Figure 2A). The last set was also used for the clustering methods comparison with r-weighted rotational averages (Figure S4D). Except when noted, 8 structures were used and 100 particles were generated for each structure. For each dataset, the corresponding ground truth classes contained particles obtained from one reference structure.

Particles were rotationally averaged around the normals (Figure S3B) and the cross-correlation between all particle pairs was calculated, to be used as the clustering distance. K-means and hierarchical classifications were preceded by principal component analysis where the eigenvectors corresponding to the eigenvalues that contributed to 95% of the variability were kept. Each K-means clustering was repeated 200 times and the mean and the standard deviations were calculated.

Classifications were evaluated by comparing them with the ground truth using the following three metrics. Fowlkes-Mallows [27] and the Variation of information [28] were implemented in Pyto package as previously reported [16]. To calculate F_1_, for each evaluated class we determined the number of particles belonging to the reference classes. The reference class that contributed the highest number of particles was then associated to the evaluated class, thus defining a mathematical map from the evaluated to the reference classes. We note that this mapping does not have to be one-to-one or onto. Nevertheless, it allows us to calculate the following properties for each reference class *i*:

- True positives (*TP*_*i*_): Number of elements of all evaluated classes associated with reference class *i* that belong to the class *i*
- False positives (*FP*_*i*_): Number of elements of all evaluated classes associated with reference class *i* that do not belong to the class *i*
- False negatives (*FN*_*i*_): Number of elements of all evaluated classes not associated with reference class *i*, that belong to the reference class *i*

Now we can define precision (*P*), recall (*R*) and the F_1_ measure:

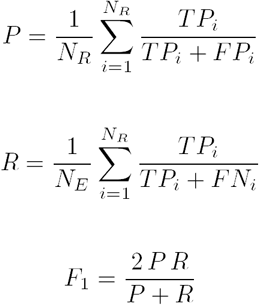

where *N*_*R*_ and *N*_*E*_ and the number of reference and evaluated classes, respectively. In this way, persistence and recall show the expected behavior, that is precision increases and recall decreases with the number of classes (Figure S4A and B).

### 3D classification and refinement

3D classification and refinement steps were performed in Relion [22]. During the refinement, particle half-datasets were processed independently according to the “gold-standard” procedure, as implemented in Relion. The resolution was determined by Fourier shell correlation at the FSC = 0.143 criterion. The constrained refinement and 3D classification were carried out by initially aligning particles according to the direction of their normal vectors. The alignment was then optimized by allowing small changes in the two normal vector angles and small spatial displacements. The alignment around the third angle (around the normal vector) was not constrained to explore the entire angular range, except when a high symmetry is used (typically C10). Specifically, we used the two angles defining the normals to the membrane to set the prior values for angles *tilt* and *psi* in Relion particle star files and specified small values (3.66) for the standard deviations of these two angles as the refine command options. Except for the validation of particle picking, the initial reference was obtained by aligning all particles according to the two angles determined from normals and randomizing the third angle (around the normal direction) to remove the missing wedge, that is no external reference was used.

### Processing of microsomal complexes

The processing work-flow is schematically shown in Figure S6.

55 out of 210 the tomograms previously obtained from canine pancreatic microsomes [29] were used in this study. From these tomograms (1.048 nm pixel size), 122 microsomal membranes were segmented using the automated procedure TomoSegMemTV [63]. For tracing of densities, persistence simplification and particle picking, tomograms were smoothed by Gaussian filtering at σ = 2 / 0.8 pixels (for the cytoplasmic / lumenal sides). Density tracing and the topological simplification were performed in 3D by our procedure as described above. To account for different quality of tomograms, the persistence threshold was set so that the density of vertices (minima) on all microsomal membrane was 0.0025 nm^−3^.

To localize cytoplasmic / lumenal particles, we selected vertices that were located at 25-75 nm / 3-15 nm Euclidean and 25-50 nm / 3-30 nm geodesic distance (length of the shortest path composed of arcs) from the membrane and had up to 2.5 sinuosity (ratio of geodesic and euclidean distances). For each selected vertex, the closest connected membrane vertex was detected, and these membrane vertices were chosen to represent particles, resulting in 64 000 and 62 000 cytosolic and lumenal particles, respectively. A particle exclusion distance of 5 nm was imposed. These particle positions were used to reconstruct particles in Imod [51] at a pixel size 0.524 / 0.262 nm and with a box size 120 / 160 pixels (for the cytoplasmic / lumenal sides).

Further processing was done essentially in the same way for the cytoplasmic and lumenal particles, as follows. Particle position and normal vectors were optimized using Relion by the constrained refinement with C10 symmetry. They were rotationally averaged, and the general classification was performed using the AP clustering. The initial preference was set to −10, to prevent getting a large number of classes The resulting classes were visually inspected to select the cytosolic class showing the best cytoplasmic and lumenal densities, and the lumenal class showing the best lumenal density. These classes were averaged by constrained refinement to yield densities to be used as initial references for the further processing.

Unless noted otherwise, the masks used for the AP and for the following 3D classification and refinement steps were cylindrical, for cytoplasmic particles 40×120 pixels (radius × height) containing both cytosolic and lumenal space, and for lumenal particles 25×110 pixels, containing lumenal and only little cytoplasmic space just above the membrane.

We then performed three rounds of 3D classification. False positive particles were removed by the first 3D classification round, with constrained alignment, using all cytosolic / lumenal particles and starting from the previously obtained initial references. In the bulk cleaning variant, all particles were classified together and the best class (resembling the initial reference the most) was selected for further processing (2600 cytosolic and 21 000 lumenal particles). In the AP cleaning variant, each affinity propagation class was subjected to 3D classification and the best classes were selected (7100 cytosolic particles from 15 affinity propagation classes). The second 3D classification round was focused on the opposite sides of the particles (lumenal / cytosolic) and the third round on the smaller regions. Both 2nd and 3rd classification steps were performed without alignment, using the alignment parameters from the first 3D classification round (the masks used are shown in Figure 3 E, F). Specifically, for the cytosolic particles, the second 3D classification was focused on the ER lumen and generated ribosome class bound to different translocon species (fully assembled translocon: 3400 particles; partially assembled: 1800 particles), while the third round of 3D classification focused on the cytosolic face of the ER membrane to separate translocon-bound ribosome (1064 particles) from translocon-bound large ribosomal subunits (873 particles) (Figure 3 E). For the lumenal particles, ribosome-bound (1800 particles) and the ribosome-free translocon (11 000 particles) classes were generated during the second 3D classification step focused on the cytosolic side, while the third 3D classification round focused on the ER lumen yielded two classes representing different ribosome-free states (separate OST complexes and full translocon complexes, 2200 and 8600 particles, respectively) (Figure 3F). An exclusion distance of 15 nm was imposed in order to remove overlapping particles, likely originating from translation shifts during the alignment steps. The final classes were averaged by the constrained refinement, post-processed and the FSC curves were generated (Figure S5).

### Validation of density tracing and particle picking on microsomal membranes

For this validation, we analyzed 6 out of 55 microsomal tomograms that we used before. The Morse tracing and picking was done in the same way as before. For local density detection, tomograms were low-pass Gaussian filtered at 5 nm and density maxima were chosen. Template matching was done using Pytom software [64], our ribosome-free translocon complex average (Figure 3F) low-pass filtered at 2.5 nm served as the template and the angular increment was 12.85°. In all cases, particles were picked on the lumenal side, up to 20 nm from the membrane, at 1.05 nm pixel size, yielding 13 000 - 17 000 particles. Further processing was done with Relion at 0.26 nm pixel size. All classes were first subjected to the constrained refinement with C10 symmetry and then 3D classified into five classes with constrained alignment, using the same ribosome-free translocon complex, filtered at 3.0 nm, as the initial reference.

### Detection and classification in situ

Twelve tomograms of intact P19 cells that contained ER were used here. Membranes were segmented with TomoSegMemTV [63] without any manual intervention. Consequently, the lumenal and cytosolic sides were not distinguished in the subsequent analysis. Tomograms were low-pass Gaussian filtered at *σ* = 0.8 pixels, the density was traced using our procedure, vertex density on membranes was 0.0035 nm^−3^ (all at 1.368 nm pixel size). Particles were picked in the region 3-10 nm away from membranes, having geodesic distance to the membrane of 3-30 nm. The exclusion distance was 5 nm to allow a rather comprehensive picking of small complexes. All together, 172 000 particle subvolumes were reconstructed at 0.342 nm pixel size, with box size of 160 pixels.

Particle position and normal vectors were optimized using Relion by the constrained refinement with C10 symmetry. They were rotationally averaged and the dot product between them was used as the similarity measure for the subsequent three rounds of AP classification, the initial preference parameter was −6.

The first AP classification (at 0.684 nm pixel size) served to discard classes that did showed two, or did not show any well resolved membrane (34 000 particles), and to group other classes into datasets defined by the following features: large densities on the particle side of the membrane (28 000 particles), large densities on the side opposite from the particles (25 000 particles) and the small densities (85 000 particles) (Figures S8, S9). In the second round (at 0.342 nm pixel size), these three datasets were AP-classified separately. Classes were were visually evaluated and grouped for further processing based on desired features (ribosome-containing and other classes from the large densities on the particle side dataset, and ribosome containing classes from the large densities on the opposite side dataset), and some classes were discarded. The selected particles were subjected to the constrained refinement with C10 symmetry by Relion, to optimize their positions and normals. The mask used for the first two AP rounds contained membranes and the region on both sides of the membrane. The third round of AP-classification (at 0.342 nm pixel size) was performed using a mask that did not contain the membrane, to improve the position and orientation of particles. Selected classes were subjected to constrained refinement, and in some cases also to 3D classification, by Relion. The third AP classification was not needed for classes containing ribosomes, from the large density on the particle side dataset.

Averages of selected classes were obtained by constrained refinement using Relion. In some cases the refinement was preceded by 3D classification. No symmetry was used except that C4 was used for the class tentatively identified as the IP3 receptor. No external structures were used, all initial references were provided from the AP classes. Final numbers of averaged particles are: 875 ribosomes, 180 ribosome-associated translocons, 63 ribosome-free translocons, 41 IP3 receptors, and 108 and 177 PLC complexes. Overlapping particles were removed. All masks used were cylindrical.

### Spatial distribution functions

We implemented the following spatial distribution functions. The nearest neighbor distribution function *G(r)* of a particle set is defined as a probability that the nearest neighbor of a particle is found at a distance ≤ *r*. The spherical contact distribution *F(r)* is a probability that the closest particle from an arbitrary point is found at a distance ≤ *r*. Consequently, *G(r)* primarily describes the organization within particle clusters, while *F(r)* the empty space. The J-function is a combination of the two:

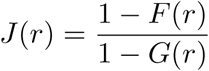

Ripley’s L function is calculated considering the region within which the particles are detected (particle region), which can have an arbitrary shape, as follows:

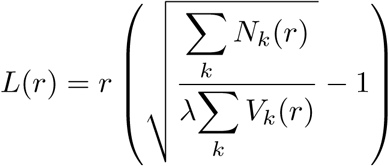

where *λ* is the overall concentration of particles, *N*_*k*_(*r*) is number of points (particles) within a distance ≤*r* to point *k*, and *V*_*k*_(*r*) is the volume of the intersection of the particle region and a sphere of radius *r* centered on the point *k* [35, 36]. The bivariate versions of functions *G(r)* and *L(r)* characterize the colocalization of two particle sets. They differ from the univariate functions specified above in that the distances are calculated from particles of one set to the particles of the other set.

We calculated Ripley’s L function for particles on each microsome and obtained the mean. For the determination of the statistical significance of Ripley’s L function, we generated multiple random distributions (10 for each microsome, that is ≈1200 total) that had the same number of particles and were located within the same spatial region as the particle set. Random points were given real particle size and were not allowed to overlap among the same class, effectively imposing an exclusion distance within a class. The envelope within which 95% of the curves were located was then used to determine whether the distribution of the particle set was significantly different from the random distribution (at the p*<*0.05 significance level).

### General software methods

The software developed for the work presented here was developed in the object-oriented manner in Python as PySeg package. The package contains installation and usage instructions, examples on real biological data and more than 66 000 lines (instructions and examples excluded). It is open-source and available upon demand.

PySeg uses Pyto package [16], Numpy package, surface meshes were stored using VTK [65] and graphs are plotted using *matplotlib* library [66]. We parallelized (shared memory model) some of the most intensive operations in order to provide a software package able to effectively process big datasets with many particles and tomograms.

For visualization, Paraview [67] and the UCSF Chimera package from the Computer Graphics Laboratory, University of California, San Francisco [68] software packages were used.

All computations were done on Linux clusters at the computer center of the Max Planck Institute of Biochemistry.

## Acknowledgements

We would like to thank Florian Beck for useful discussions and Gabriela J. Greif for critical reading of the manuscript. A.M.-S. is the recipient of a postdoctoral fellowship from the Séneca Foundation.

## Author Contributions

AM-S and VL conceived and designed the research. AM-S designed and implemented the software; ZK, UL and SC acquired original tomograms; SP provided expertise related to the previously recorded tomograms; AM-S, JMzAB and VL analyzed the data; WB provided resources and acquired funding; VL supervised research; AM-S and VL wrote the manuscript. All authors edited the manuscript.

## Additional information

### Accession codes

EM densities have been deposited in the EMDataBank obtained from cytosolic particles: ribosome bound to the fully assembled (EMD-0074) and partial translocon complex (EMD-0084), the large ribosomal subunit (EMD-0075), and obtained from lumenal particles: the ribosome-translocon complex (EMD-0085), the ribosome-free fully assembled translocon (EMD-0086) and the non-translocon associated OST complex (EMD-0087).

## References

[1] Santos, R. et al. A comprehensive map of molecular drug targets. Nature reviews. Drug discovery 16, 19–34 (2017).

[2] Taylor, K. A. & Glaeser, R. M. Electron diffraction of frozen, hydrated protein crystals. Science 186, 1036–7 (1974).

[3] Dubochet, J. et al. Cryo-electron microscopy of vitrified specimens. Q Rev Biophys 21, 129–228 (1988).

[4] Lucic, V., Rigort, A. & Baumeister, W. Cryo-electron tomography: the challenge of doing structural biology in situ. J Cell Biol 202, 407–419 (2013).

[5] Oikonomou, C. M. & Jensen, G. J. Cellular electron cryotomography: Toward structural biology in situ. Annual Review of Biochemistry 86, 873–896 (2017).

[6] Medalia, O. et al. Macromolecular architecture in eukaryotic cells visualized by cryoelectron tomography. Science 298, 1209–13 (2002).

[7] Frangakis, A. S. et al. Identification of macromolecular complexes in cryoelectron tomograms of phantom cells. Proc Natl Acad Sci U S A 99, 14153–8 (2002).

[8] Ortiz, J. O., Forster, F., Kurner, J., Linaroudis, A. A. & Baumeister, W. Mapping 70S ribosomes in intact cells by cryoelectron tomography and pattern recognition. J Struct Biol 156, 334–41 (2006).

[9] Beck, M. et al. Visual proteomics of the human pathogen Leptospira interrogans. Nat Methods 6, 817–23 (2009).

[10] Rigort, A. et al. Automated segmentation of electron tomograms for a quantitative description of actin filament networks. J Struct Biol 177, 135–144 (2012).

[11] Asano, S. et al. Proteasomes. a molecular census of 26s proteasomes in intact neurons. Science 347, 439–442 (2015).

[12] Rickgauer, J. P., Grigorieff, N. & Denk, W. Single-protein detection in crowded molecular environments in cryo-em images. eLife 6, e25648 (2017).

[13] Volkmann, N. Methods for segmentation and interpretation of electron tomographic reconstructions. Methods Enzymol 483, 31–46 (2010).

[14] Fernandez, J.-J. Computational methods for electron tomography. Micron 43, 1010–1030 (2012).

[15] Chen, M. et al. Convolutional neural networks for automated annotation of cellular cryo-electron tomograms. Nature methods 14, 983–985 (2017).

[16] Lucic, V., Fernández-Busnadiego, R., Laugks, U. & Baumeister, W. Hierarchical detection and analysis of macromolecular complexes in cryo-electron tomograms using pyto software. Journal of structural biology 196, 503–514 (2016).

[17] Fernández-Busnadiego, R. et al. Quantitative analysis of the native presynaptic cytomatrix by cryoelectron tomography. J Cell Biol 188, 145–56 (2010).

[18] Fernández-Busnadiego, R. et al. Cryo-electron tomography reveals a critical role of rim1*α* in synaptic vesicle tethering. J Cell Biol 201, 725–740 (2013).

[19] Förster, F., Medalia, O., Zauberman, N., Baumeister, W. & Fass, D. Retrovirus envelope protein complex structure in situ studied by cryo-electron tomography. Proc Natl Acad Sci U S A 102, 4729–4734 (2005).

[20] Schur, F. K. et al. An atomic model of hiv-1 capsid-sp1 reveals structures regulating assembly and maturation. Science 353, 506–508 (2016).

[21] Wan, W. & Briggs, J. Cryo-electron tomography and subtomogram averaging. In Methods in enzymology, vol. 579, 329–367 (Elsevier, 2016).

[22] Bharat, T. A. & Scheres, S. H. Resolving macromolecular structures from electron cryo-tomography data using subtomogram averaging in relion. Nature protocols 11, 2054 (2016).

[23] Sousbie, T. The persistent cosmic web and its filamentary structure–i. theory and implementation. Monthly Notices of the Royal Astronomical Society 414, 350–383 (2011).

[24] Milnor, J. Morse theory, volume 51 of annals of mathematics studies. Princeton, NJ, USA (1963).

[25] Forman, R. A user’s guide to discrete morse theory. Seminaire Lotharingien de Combinatoire 48, 35pp (2002).

[26] Frey, B. J. & Dueck, D. Clustering by passing messages between data points. Science 315, 972–976 (2007).

[27] Fowlkes, E. B. & Mallows, C. L. A method for comparing two hierarchical clusterings. Journal of the American Statistical Association 78, 553–569 (1983).

[28] Meila, M. Comparing clusterings - an information based distance. Journal of Multivariate Analysis 98, 873–895 (2007).

[29] Pfeffer, S. et al. Structure of the native sec61 protein-conducting channel. Nature communications 6, 8403 (2015).

[30] Pfeffer, S. et al. Structure of the mammalian oligosaccharyl-transferase complex in the native er protein translocon. Nature communications 5, 3072 (2014).

[31] Pfeffer, S., Dudek, J., Zimmermann, R. & Förster, F. Organization of the native ribosome–translocon complex at the mammalian endoplasmic reticulum membrane. Biochimica et Biophysica Acta (BBA)-General Subjects 1860, 2122–2129 (2016).

[32] Fan, G. et al. Gating machinery of insp3r channels revealed by electron cryomicroscopy. Nature 527, 336–341 (2015).

[33] Blees, A. et al. Structure of the human mhc-i peptide-loading complex. Nature 551, 525–528 (2017).

[34] Stoyan, D. Fundamentals of point process statistics. In Baddeley, A., Gregori, P., Mateu, J., Stoica, R. & Stoyan, D. (eds.) Case studies in spatial point process modeling (Springer, 2006).

[35] Ripley, B. D. Spatial statistics (Willey-Interscience, 1981).

[36] Wiegand, T. & Moloney, K. A. Rings, circles, and null-models for point pattern analysis in ecology. Oikos 104, 209–229 (2004).

[37] Kudryashev, M. et al. The structure of the mouse serotonin 5-ht3 receptor in lipid vesicles. Structure (London, England: 1993) 24, 165–170 (2016).

[38] Xu, M., Beck, M. & Alber, F. Template-free detection of macromolecular complexes in cryo electron tomograms. Bioinformatics 27, i69–i76 (2011).

[39] Pei, L., Xu, M., Frazier, Z. & Alber, F. Simulating cryo electron tomograms of crowded cell cytoplasm for assessment of automated particle picking. BMC Bioinformatics 17, 405 (2016).

[40] Xu, M. et al. De novo structural pattern mining in cellular electron cryotomograms. Structure (London, England: 1993) (2019).

[41] Shrimal, S., Cherepanova, N. A. & Gilmore, R. DC2 and KCP2 mediate the interaction between the oligosaccharyltransferase and the ER translocon. J Cell Biol 216, 3625–3638 (2017).

[42] Braunger, K. et al. Structural basis for coupling protein transport and N-glycosylation at the mammalian endoplasmic reticulum. Science 360, 215–219 (2018).

[43] Andronov, L. et al. 3dclustervisu: 3d clustering analysis of super-resolution microscopy data by 3d voronoi tessellations. Bioinformatics (Oxford, England) 34, 3004–3012 (2018).

[44] Dunkley, P. R. et al. A rapid percoll gradient procedure for isolation of synaptosomes directly from an s1 fraction: homogeneity and morphology of subcellular fractions. Brain Res 441, 59–71 (1988).

[45] Godino, M. d. C., Torres, M. & Sánchez-Prieto, J. Cb1 receptors diminish both ca(2+) influx and glutamate release through two different mechanisms active in distinct populations of cerebrocortical nerve terminals. J Neurochem 101, 1471–1482 (2007).

[46] Koster, A. J. et al. Perspectives of molecular and cellular electron tomography. J Struct Biol 120, 276–308 (1997).

[47] Mastronarde, D. N. Automated electron microscope tomography using robust prediction of specimen movements. J Struct Biol 152, 36–51 (2005).

[48] Danev, R., Buijsse, B., Khoshouei, M., Plitzko, J. M. & Baumeister, W. Volta potential phase plate for in-focus phase contrast transmission electron microscopy. Proc Natl Acad Sci USA 111, 15635–15640 (2014).

[49] Zheng, S. Q. et al. Motioncor2: anisotropic correction of beam-induced motion for improved cryo-electron microscopy. Nature methods 14, 331–332 (2017).

[50] Grant, T. & Grigorieff, N. Measuring the optimal exposure for single particle cryo-em using a 2.6 Å reconstruction of rotavirus vp6. eLife 4, e06980 (2015).

[51] Kremer, J. R., Mastronarde, D. N. & McIntosh, J. R. Computer visualization of three-dimensional image data using imod. J Struct Biol 116, 71–76 (1996).

[52] Nash, C. & Sen, S. Topology and geometry for physicists (Academic Press, Harcourt Brace Jovanovic, London, 1990).

[53] Peixoto, T. P. The graph-tool python library. figshare (2014).

[54] Pedregosa, F. et al. Scikit-learn: Machine learning in Python. Journal of Machine Learning Research 12, 2825–2830 (2011).

[55] Meyerson, J. R. et al. Structural mechanism of glutamate receptor activation and desensitization. Nature 514, 328–334 (2014).

[56] Karakas, E. & Furukawa, H. Crystal structure of a heterotetrameric nmda receptor ion channel. Science (New York, N.Y.) 344, 992–997 (2014).

[57] Herguedas, B. et al. Structure and organization of heteromeric ampa-type glutamate receptors. Science (New York, N.Y.) 352, aad3873 (2016).

[58] Wu, J. et al. Structure of the voltage-gated calcium channel ca(v)1.1 at 3.6?Å resolution. Nature 537, 191–196 (2016).

[59] Morales-Perez, C. L., Noviello, C. M. & Hibbs, R. E. X-ray structure of the human ?4?2 nicotinic receptor. Nature 538, 411–415 (2016).

[60] Tao, X., Hite, R. K. & MacKinnon, R. Cryo-em structure of the open high-conductance ca, javax.xml.bind.jaxbelement@741fc2e3, -activated k, javax.xml.bind.jaxbelement@4349c43b, channel. Nature 541, 46–51 (2017).

[61] Park, E., Campbell, E. B. & MacKinnon, R. Structure of a clc chloride ion channel by cryo-electron microscopy. Nature 541, 500–505 (2017).

[62] Zhang, Y. et al. Cryo-em structure of the activated glp-1 receptor in complex with a g protein. Nature 546, 248–253 (2017).

[63] Martinez-Sanchez, A., Garcia, I., Asano, S., Lucic, V. & Fernandez, J.-J. Robust membrane detection based on tensor voting for electron tomography. J Struct Biol 186, 49–61 (2014).

[64] Hrabe, T. et al. Pytom: a python-based toolbox for localization of macromolecules in cryo-electron tomograms and subtomogram analysis. Journal of structural biology 178, 177–188 (2012).

[65] [65] Schroeder, W. J., Lorensen, B. & Martin, K. The visualization toolkit: an object-oriented approach to 3D graphics (Kitware, 2004).

[66] Hunter, J. D. Matplotlib: A 2d graphics environment. Comput. Sci. Eng. 9, 90–95 (2007).

[67] Ayachit, U. The paraview guide: a parallel visualization application (2015).

[68] Pettersen, E. F. et al. Ucsf chimera, a visualization system for exploratory research and analysis. J Comput Chem 25, 1605–12 (2004).

