## Supplemental data for "Template-free detection and classification of heterogeneous membrane-bound complexes in cryo-electron tomograms"

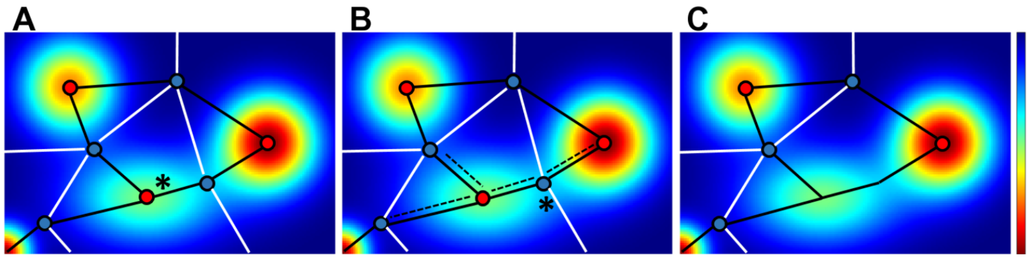


Figure S1: Simplification by topological persistence in 2D. A The initial Morse complex is shown on the density map. The minimum to be removed by the simplification is labeled by an asterisk. B Arcs that are to be modified are indicated by dashed lines and the saddle point that is to be removed is indicated by an asterisk. C Morse complex after the simplification. On all panels minima are shown by red and saddle points as blue points, arcs are shown as black lines and the white lines denote borders between ascending 2-manifolds.

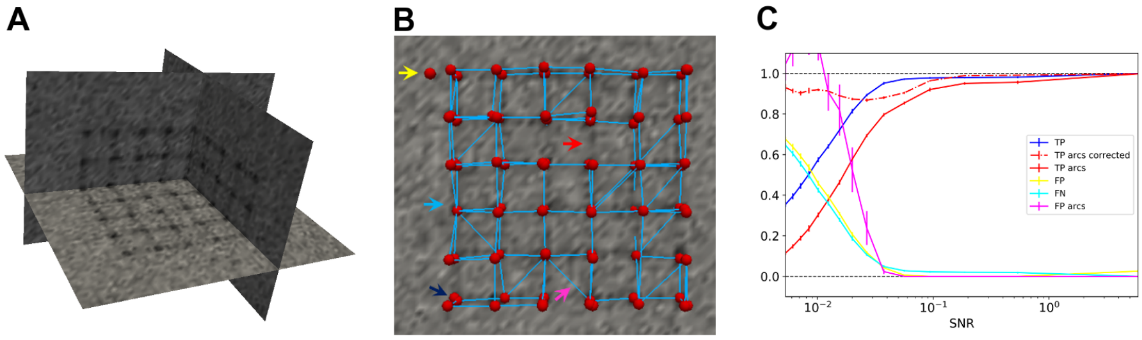


Figure S2: Validation of the density tracing and the simplification on phantom data. A Tomographic slices of the phantom data set. B Detected minima and arcs at SNR = 0.025 in the graph representation superposed on a slice of the simulated tomogram. Red spheres denote detected minima (vertices) and blue lines detected arcs (edges). The yellow arrow points to a FP minimum, blue to a FN minimum, dark blue to a double minimum, pink to a FP arc and red to a FN arc. C Normalized number of TP, FP and FN minima (labeled as TP, FP and FN), and TP and FP for arcs (mean ± std, N=10 simulations per SNR).

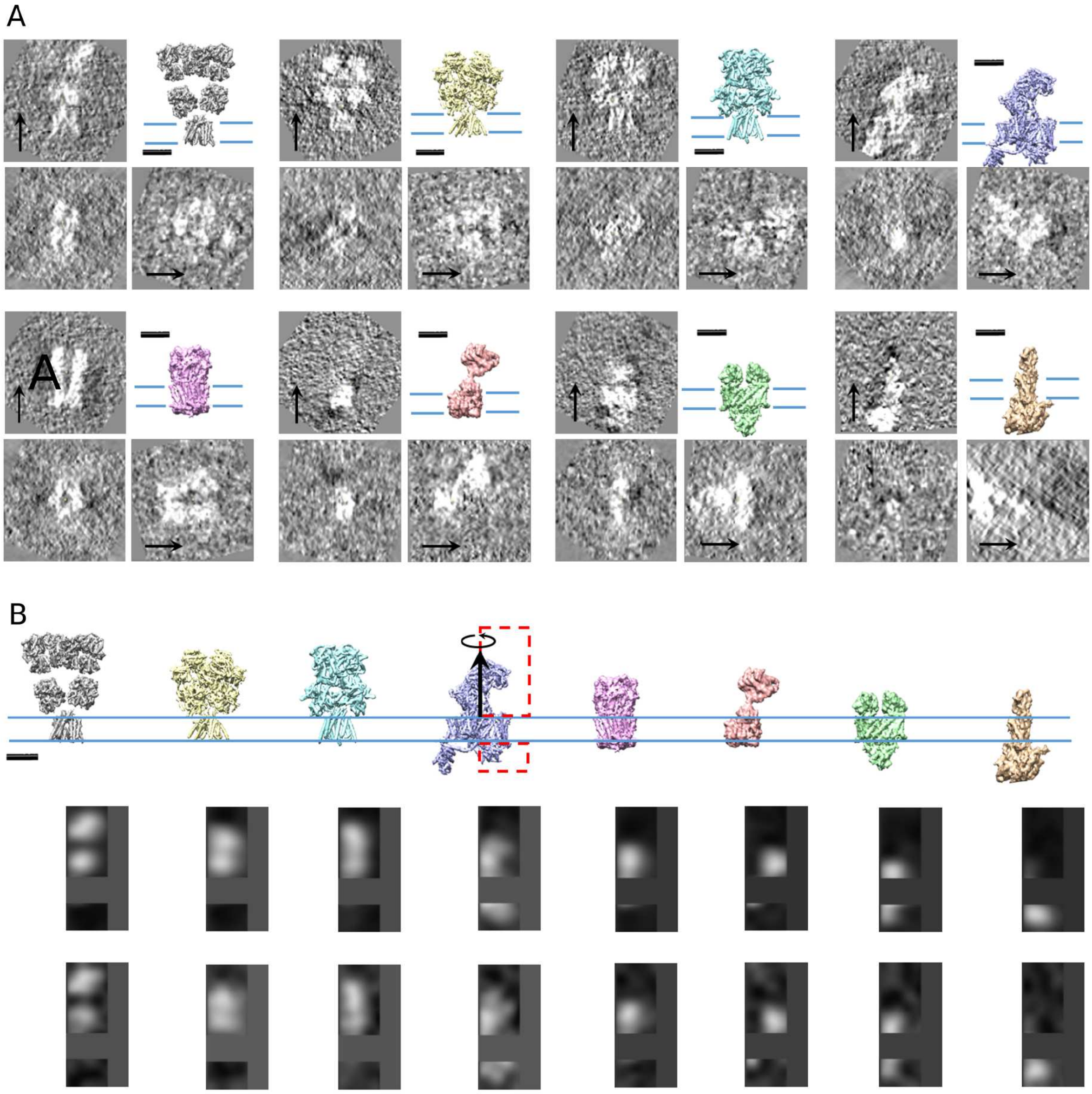


Figure S3: Particles used for the valuation of AP classification. A High resolution structures and particles used to make the test data (SNR = 0.05). B High resolution structures used to make test data sets (top row), corresponding AP class averages of the test dataset used for the comparison of the clustering methods (middle row) and AP class exemplars obtained using the same dataset (bottom row). Scale bars 5 nm.

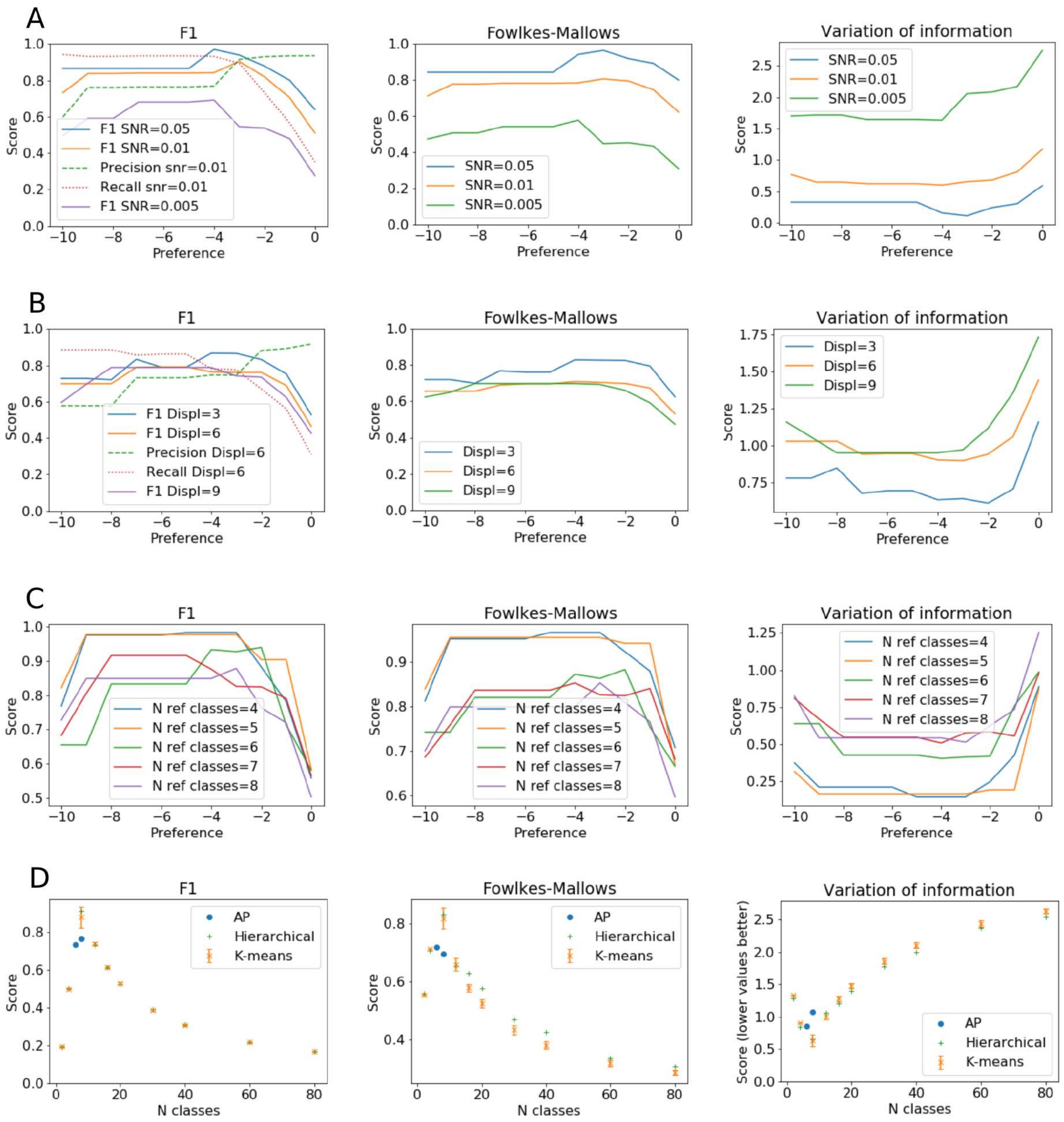


Figure S4: Evaluation of AP classification. A Evaluation of classifications obtained for different SNR's and their dependence on the preference parameter. B Evaluation of classifications obtained for different displacement ranges along the membrane and their dependence on the preference parameter. C Evaluation of classifications obtained for different number of reference classes and their dependence on the preference parameter. D Comparison of the AP with K-means and hierarchical clusterings, where particles were rotationally averaged with r-weighting. The AP clustering results are shown for the preference parameter taking all integer values between -9 and -3 (note that several data points overlap). For K-means clustering the data points denote means and the error bars the standard deviation obtained from 200 simulations. Fowlkes-Mallows (left), Variation of information (middle) and $F_1$ measures (right). Note that higher values of the Fowlkes-Mallows and $F_1$ measures, but lower values of the Variation of information signify better agreement. Note that the number of classes increases with the preference parameter.

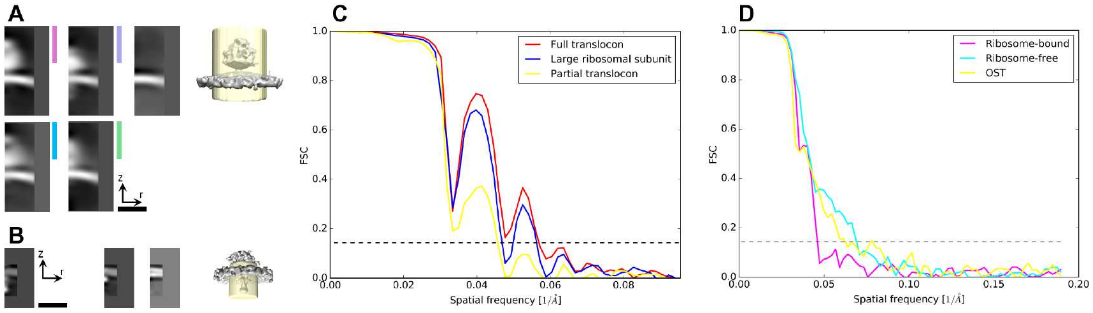


Figure S5: Processing of microsome-attached complexes. A Affinity propagation classification of cytosolic particles (ribosomes). 2D class averages are shown on the left side, the four color bar labeled images are from the same classes as those shown in the same colors on Figure 3C, D. The class shown in the upper left corner was used to generate the initial reference for the subsequent 3D classification. 3D class average obtained without alignment is shown on the right, the mask is shown in transparent yellow. B Affinity propagation classes of lumenal particles (translocons). 2D class averages are shown on the left. The class shown on the right was used as the initial reference for the subsequent 3D classification. 3D class average obtained without alignment is shown on the right, the mask is shown in transparent yellow. C FSC plots of the 3D refined ribosomal class averages obtained using the AP cleaning variant, shown on Figure 3E (colors match). D FSC plots of the 3D refined translocon class averages shown on Figure 3F (colors match). FSC plots in C and D were obtained by the AP cleaning variant. Scale bars 20 nm.

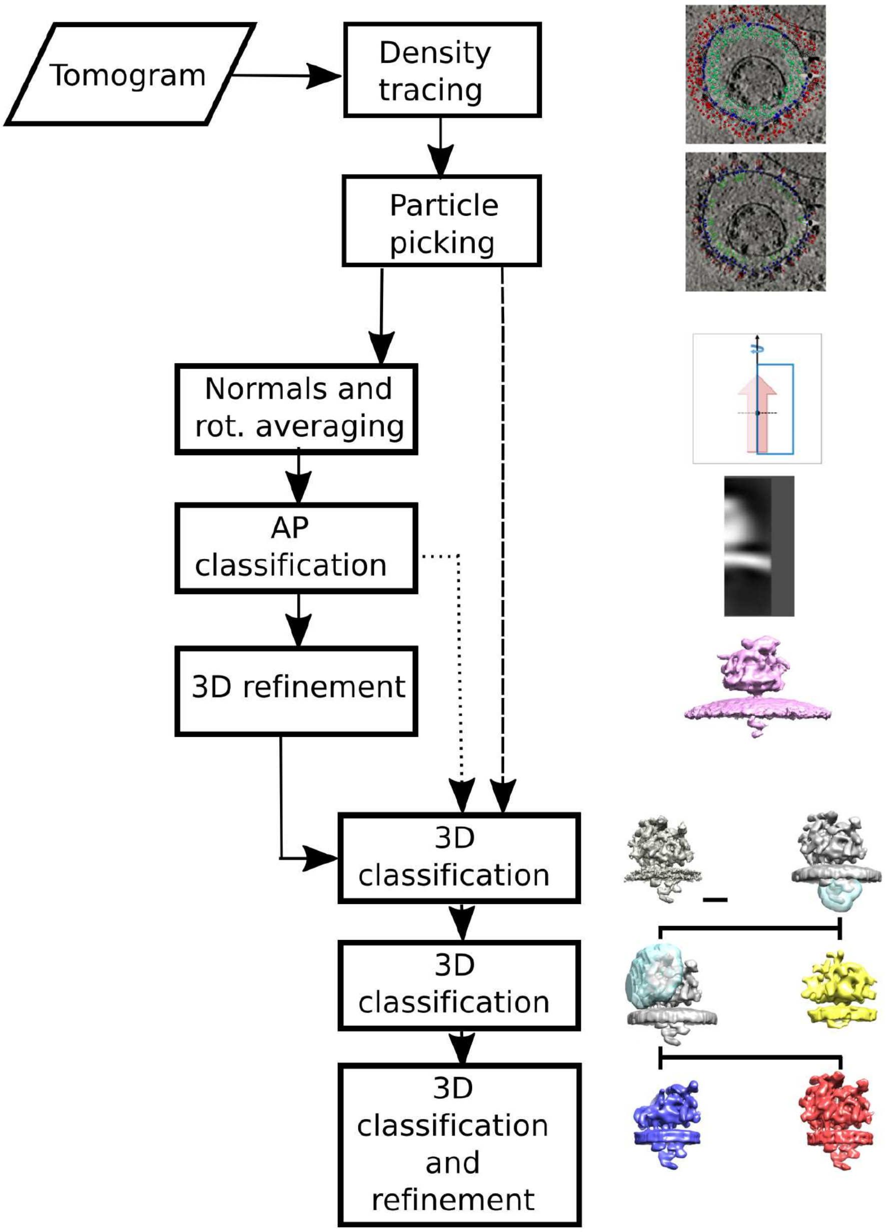


Figure S6: Flowchart of the processing steps used for microsome-attached complexes. Rectangles show the computational steps. Solid lines and the dotted line together show the processing flow that includes the AP cleaning variant of the first 3D classification step. The dashed line together with the solid lines show the bulk cleaning variant. Images on right side graphically represent the results of the corresponding .processing steps.

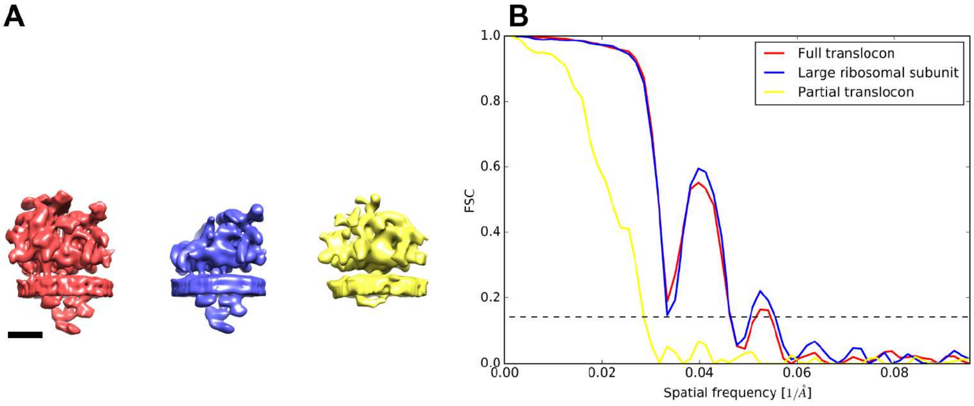


Figure S7: Processing lumenal particles using the bulk cleaning variant. A 3D densities of fully assembled (left) and partial translocon complex (right), as well as the large ribosomal subunit bound to the fully assembled translocon (middle). B FSC plots of the classes shown in A. Colors correspond to the densities shown in Figure 3E. Scale bar 10 nm.

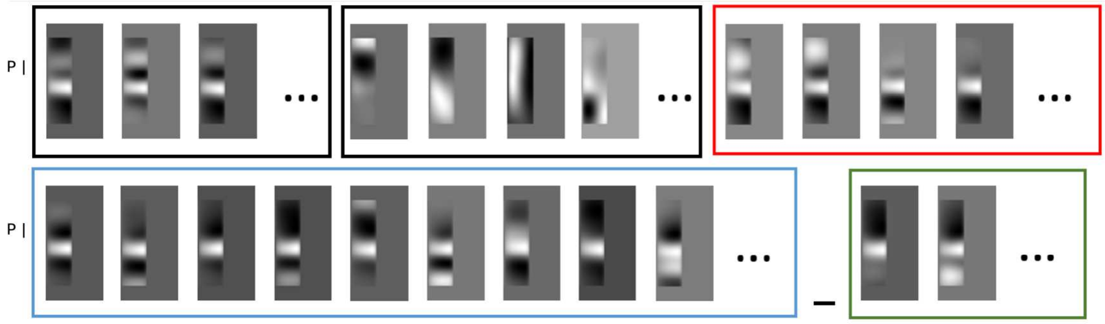


Figure S8: Examples of class averages obtained after the first AP classification round. Vertical lines on the left labeled with “P” show the level where particles were picked. The two groups boxed in black show classes containing double membranes (left) and classes not containing a clear membrane (middle). The group boxed in red is shown in Figure S9A, blue in Figure S9B and green in Figure S9C. Scale bar 10 nm.

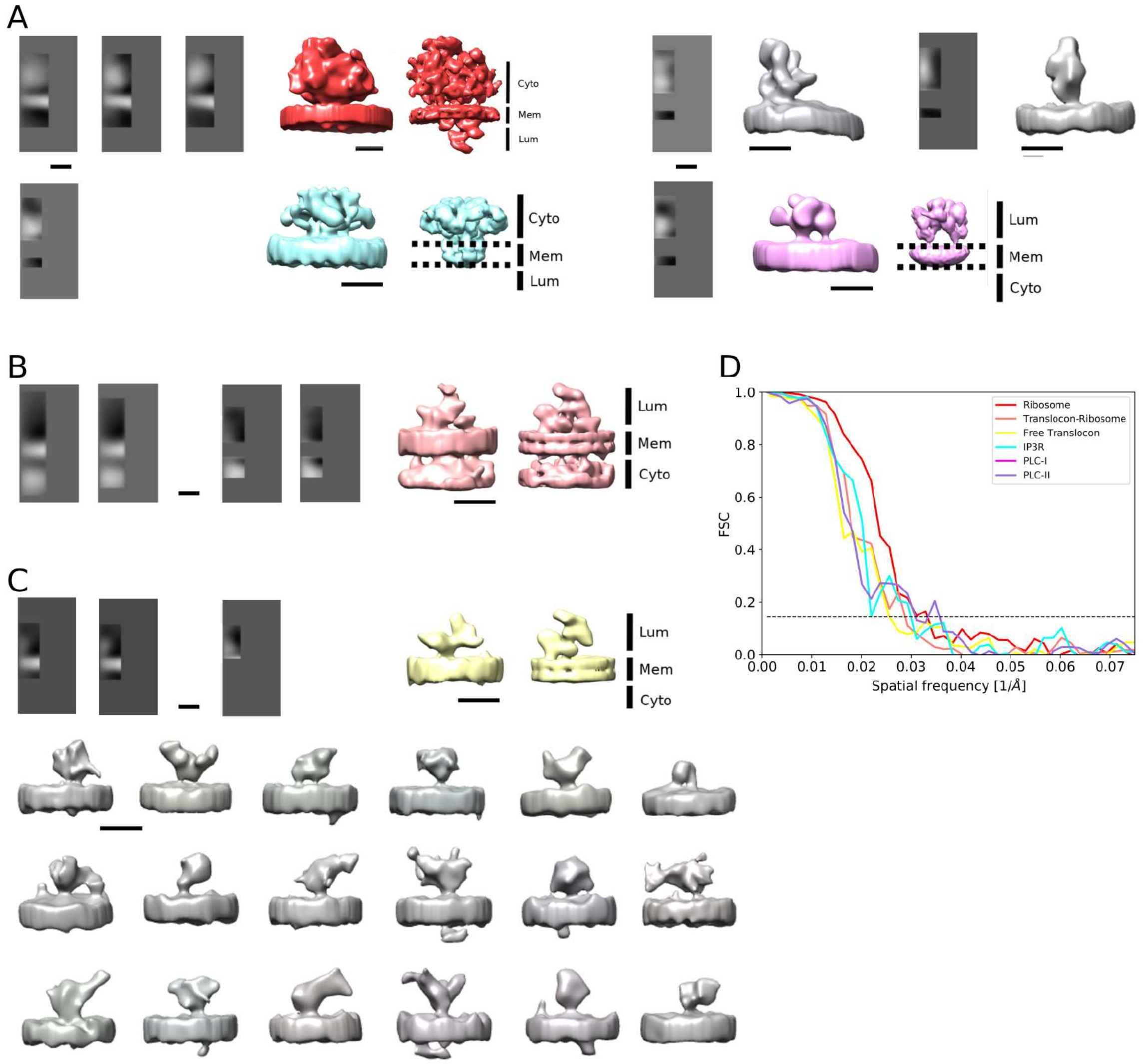


Figure S9: Classification and refinement of in situ complexes. A The third AP classification and refinement of the dataset featuring large densities on the particle side of the membrane. Top left: ribosomes (AP averages, in situ 3D average and the 3D average obtained from microsomes). Top right: two unidentified structures (AP average, in situ 3D average). Bottom left: Putative IP3 receptor (AP average, in situ 3D average and EMD-6369). Bottom right: putative PLC (AP average, in situ 3D average and EMD-3905). B Dataset featuring large densities on the side opposite of the particle. Ribosome-bound translocon (the second and the third AP classifications, 3D in situ average and the 3D average obtained from microsomes). C Dataset showing small densities. Top row: ribosome-free translocon (the second and the third AP classifications, 3D in situ average and the 3D average obtained from microsomes). Below: a gallery showing other average densities from the same dataset. In situ 3D averages are also shown on Figure 4, 3D average obtained from microsomes are also shown on Figure 3). D FSC plots of the average densities shown in color. All scale bars 10 nm.

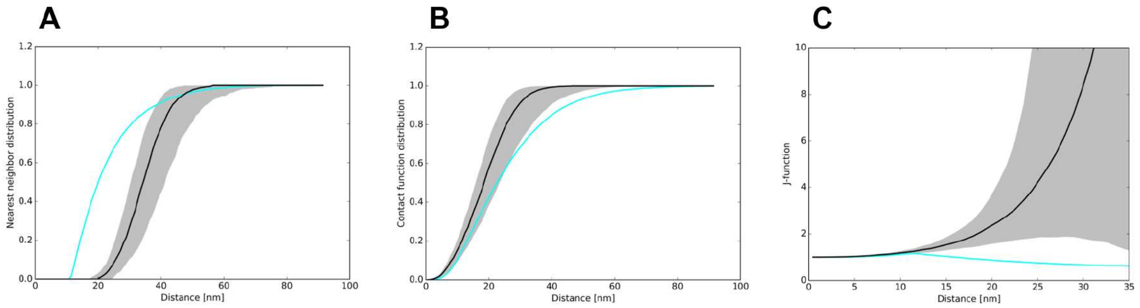


Figure S10: Clustering of ribosome-free full translocon by the first order distribution functions. A Nearest neighborhood distribution. B Contact distribution. C J-function.

Video 1: Detection of microsome-attached complexes and localization of lumenal particles. Density minima are shown as small spheres (red: cytosolic; blue membrane; green lumenal), arcs as grey lines and green arrows denote membrane normal vectors.

Video 2: Localization of microsome-attached complexes. Ribosomes derived from the cytosolic particles are shown in red, ribosme-free fully assembled translocon complexes in blue and the non-translocon associated OST complexes in yellow.
